# Cytolytic memory CD4^+^ T cell clonotypes are expanded during *Plasmodium falciparum* infection

**DOI:** 10.1101/2021.07.21.453277

**Authors:** Raquel Furtado, Fabien Delahaye, Jinghang Zhang, Joowhan Sung, Paul Karell, Ryung S. Kim, Sophie Caillat-Zucman, Li Liang, Philip Felgner, Andy Bauleni, Syze Gama, Andrea Buchwald, Terrie Taylor, Karl Seydel, Miriam Laufer, Johanna P. Daily, Grégoire Lauvau

**Affiliations:** Department of Microbiology and Immunology; Department of Genetics; Department of Medicine; Department of Epidemiology and Population Health, Albert Einstein College of Medicine, Bronx, New York, USA 10461; Université de Paris, AP-HP, Hôpital Saint-Louis, Laboratoire d’Immunologie et Histocompatiblité, INSERM UMR976, 75010 Paris, France; Department of Physiology and Biophysics, School of Medicine, University of California, Irvine, California, USA 92697.; Malaria Alert Centre, Kamuzu University of Health Sciences, Blantyre, Malawi; Blantyre Malaria Project, Kamuzu University of Health Sciences, Blantyre, Malawi; 9Center for Vaccine Development and Global Health, University of Maryland School of Medicine, Baltimore, Maryland, USA 21201; Department of Osteopathic Medical Specialties, Michigan State University, East Lansing, Michigan, USA 48824

## Abstract

*Plasmodium falciparum* (*Pf*) malaria causes high rates of morbidity and mortality and lacks a sufficiently effective vaccine. Clinical immunity develops in residents of malaria endemic regions which confers reduced clinical symptoms during infection and protection against severe disease. We hypothesized that understanding the immune mechanisms of clinical immunity could inform vaccine design to improve efficacy. We compared the peripheral blood cellular and humoral immune responses during a mild episode of *Pf* malaria infection. Participants were classified as either clinically susceptible or clinically protected, based on the number of recurrent clinical infections over an 18-month longitudinal study in a malaria endemic region in Malawi. Susceptible participants had three or more recurrent clinical episodes while clinically immune individuals had one or none. Protected participants exhibited higher plasma immunoglobulin G (IgG) breadth and titers against *Pf* antigens, and greater antibody (Ab)-dependent *Pf* opsonization compared to susceptible participants. Using high dimensional mass cytometry (CyTOF), spectral flow cytometry and single-cell transcriptomic analyses, we identified expanded memory CD4^+^ T cell clones sharing identical T cell receptor clonotypes in the blood of protected participants during malaria infection. These cells express a strong cytolytic T helper 1 effector program with transcripts encoding granzymes (A, B, H, M), granulysin, NKG7 and the Zeb2 master transcriptional regulator of terminally differentiated effector T cells. Memory CD4^+^ T cells expressing Zeb2^+^ were CD39^hi^TIGIT^hi^ and expressed multiple chemotactic and checkpoint inhibitory receptors, although the cellular levels of several of these receptors were reduced in protected compared to susceptible individuals. We propose that clonally expanded Zeb2^+^ cytolytic memory CD4^+^ Th1 cells could represent essential contributors to clinical immunity against *Pf* malaria.

**One Sentence Summary:** A population of cytolytic memory CD4^+^ T cells is clonally expanded in patients with *Plasmodium falciparum* malaria and has reduced chemotactic and inhibitory receptor expression in patients with naturally acquired clinical malaria immunity.

## INTRODUCTION

*Plasmodium falciparum* infection remains highly prevalent with an estimated 229 million cases of malaria worldwide and an estimated 409,000 deaths in 2019 ^1, 2^. Malaria infection is also associated with significant morbidity including decreased cognitive function and anemia in school aged children ^3–5^. The *P. falciparum* (*Pf*) parasite evades host immunity by preventing optimal T and B cell responses, and by subverting effective vaccination through a variety of mechanisms ^6–8^. These include generating immense antigenic diversity, also through distinct stages of development ^9^, promoting inflammation ^10–12^ and inducing multiple immune suppressive mechanisms ^13–17^. Nevertheless, residents of highly malaria endemic regions with high transmission develop clinical immunity, a host state that protects them from severe illness and death despite infection. Patients with clinical immunity have mild to no clinical symptoms and have low parasite burdens that are often submicroscopic during infection ^18–20^. Protection from severe disease and the lack of symptoms during infection appears to be long lasting if residents remain in endemic regions ^21, 22^. The most developed vaccine candidate RTS,S/AS01, modestly reduces the number of clinical malaria episodes by 46% and severe malaria by 36% ^23^. In addition, the efficacy of RTS,S/AS01 wanes over time, with only 2.5% efficacy four years after vaccination, with some vaccinees having a rebound in the number of clinical malaria episodes during the fifth year ^24, 25^. Thus, a more effective and long-lasting vaccine is still needed. Studies of naturally acquired clinical immunity may shed light on immune mechanisms that could be leveraged in the development of a better vaccine. How natural protection is achieved remains poorly understood but could serve as a steppingstone to uncover novel immune mechanisms relevant to host immunity against *P. falciparum*.

Both innate and adaptive responses have been associated with host protection against *P. falciparum* infection and clinical disease, and long-term immunological memory relies on parasite-specific CD4^+^ T, CD8^+^ T and B lymphocytes. In individuals vaccinated with irradiated sporozoites and subsequently challenged with homologous sporozoite parasites in human and animal studies, polyfunctional memory CD8^+^ and possibly CD4^+^ T cells help mediate robust liver stage protection ^26–32^. The liver stage of infection represents a *Pf* parasite population bottleneck, where there is limited parasite genetic variation, and effective protection could possibly be achieved. It is known that IFNγ and cytolysis of infected hepatocytes are involved, however, the exact effector mechanism that are needed and the sequences of events leading to host protection are still being debated. Whether any these mechanisms are implicated in naturally acquired immunity also remains to be determined.

Once merozoites are released from the liver to the blood circulation, which occurs within ∼7 days in humans, parasite-specific antibodies (Abs), most notably IgG ^33–36^ but also IgM ^37, 38^, represent essential mediators of naturally acquired immunity. The production of parasite-specific protective Abs is a lengthy and complex process that may take years of endemic exposure to malaria, likely as a result of multiple negative regulatory mechanisms ^39^. Serum Abs from clinically protected patients react against a higher number of parasite antigens (Ags) and have increased titers compared to sera from susceptible individuals ^39, 40^, but which parasite Ags are targeted and whether specific functional features of targeting Abs are necessary to achieve clinical immunity is still unknown. Parasite-specific CD4^+^ T cells, especially follicular helper CD4^+^ T (TFH) cells, are required for the induction of protective Ab-secreting parasite-specific B cells. However, the development of a CD4^+^ TFH cell response that promotes effective and long-lasting parasite-specific protective Abs results from multiple competing mechanisms orchestrating CD4^+^ T cell differentiation during infection. Malaria infections, both in humans and in mouse models, drive robust T helper 1 (TH1) responses as a result of a highly inflammatory environment, in which CD4^+^ T cells upregulate the TH1 master transcriptional regulator T-bet, produce IFNγ, and have cytolytic potential ^41–43^, all of which contribute to protective antimalarial responses ^44^. These T cells also express high levels of cell-surface CXCR3 and have poor TFH cell functional characteristics ^10, 45, 46^. However, as the inflammatory response decreases in individuals developing naturally acquired immunity, less TH1 polarized CD4^+^ T cells develop and differentiate into CXCR5^+^PD-1^+^CXCR3^-^ CD4^+^ T cells which exhibit the functional phenotype of TFH cells and the ability to provide strong help to parasite-specific B cells. Highly differentiated IFNγ^+^ CD4^+^ T cells can also co-secrete the immunoregulatory cytokine IL-10, dampening the immunopathology associated with the excessive inflammatory responses driven by the parasite during acute infection ^17, 47, 48^. Thus, CD4^+^ TH cells are essential for effective protection against malaria, yet exactly how these subsets are related to each other during malaria infection, how they develop during acquisition of natural immunity and contribute to protection, is not clear. The functional features that these cells need to acquire to provide an effective host antimalarial response are mostly unknown.

In the current work, we undertook a comprehensive, unbiased systems immunology approach to define humoral and cellular immune signatures during symptomatic *Pf* infection in residents of a high transmission malaria endemic area in Malawi. Participants were classified as clinically susceptible or clinically immune based on the number of clinical malaria episodes per participant over an 18-month longitudinal study. Using these clinical classifications, we then characterized their *Pf-*specific Ab response and conducted high dimensional flow cytometry phenotyping and single cell transcriptomic analysis of their memory CD4^+^ T cells to identify signatures associated with clinical immunity against malaria. Our results show that participants with clinical immunity have greater parasite-specific Ab breadth, titers and merozoite opsonization compared to susceptible subjects. Importantly, we reveal the existence of a population of memory CD4^+^ T cells expressing a robust cytolytic signature and the Zeb2 master regulator of terminally differentiated effector cells in clinically immune participants. These cells are clonally expanded during *Pf* infection and share identical TCR clonotypes and may play a key role in clinical immunity.

## RESULTS

### Longitudinal study of *Pf-*infected patients in Chikwawa District, Malawi

To identify immune signatures associated with clinical immunity against malaria, we conducted an 18-month longitudinal study of pediatric and adult patients residing in a rural region of southern Malawi in the Chikwawa District. This is a *Pf* malaria endemic area with an inoculation rate of 183 infective bites/person per year, with year-round and high transmission and where older age is associated with clinical immunity ^49^. To capture a continuum of clinical immunity and compare their immune responses, we enrolled equal numbers of children, adolescents, and adult patients (Figure 1A and Figure S1A). One hundred and twenty patients, who were otherwise healthy with no chronic conditions were enrolled during an episode of symptomatic, mild malaria infection, which was confirmed by microscopy of the blood smear. Participants were then evaluated monthly and at interim study clinic visits during illness to monitor for malaria infection, record temperature and collect blood samples over the course of the study. Clinical reinfection was defined as symptoms consistent with malaria and >2,500 parasites/μl seen on microscopy. Ninety-seven participants completed the study at 18 months of follow up (Figure 1A) with age-related characteristics reported in Figure S1A. Two protected participants had sickle cell trait and were excluded from further analysis as sickle cell confers a unique protective mechanism against malaria infection ^50^.

**Figure 1.**
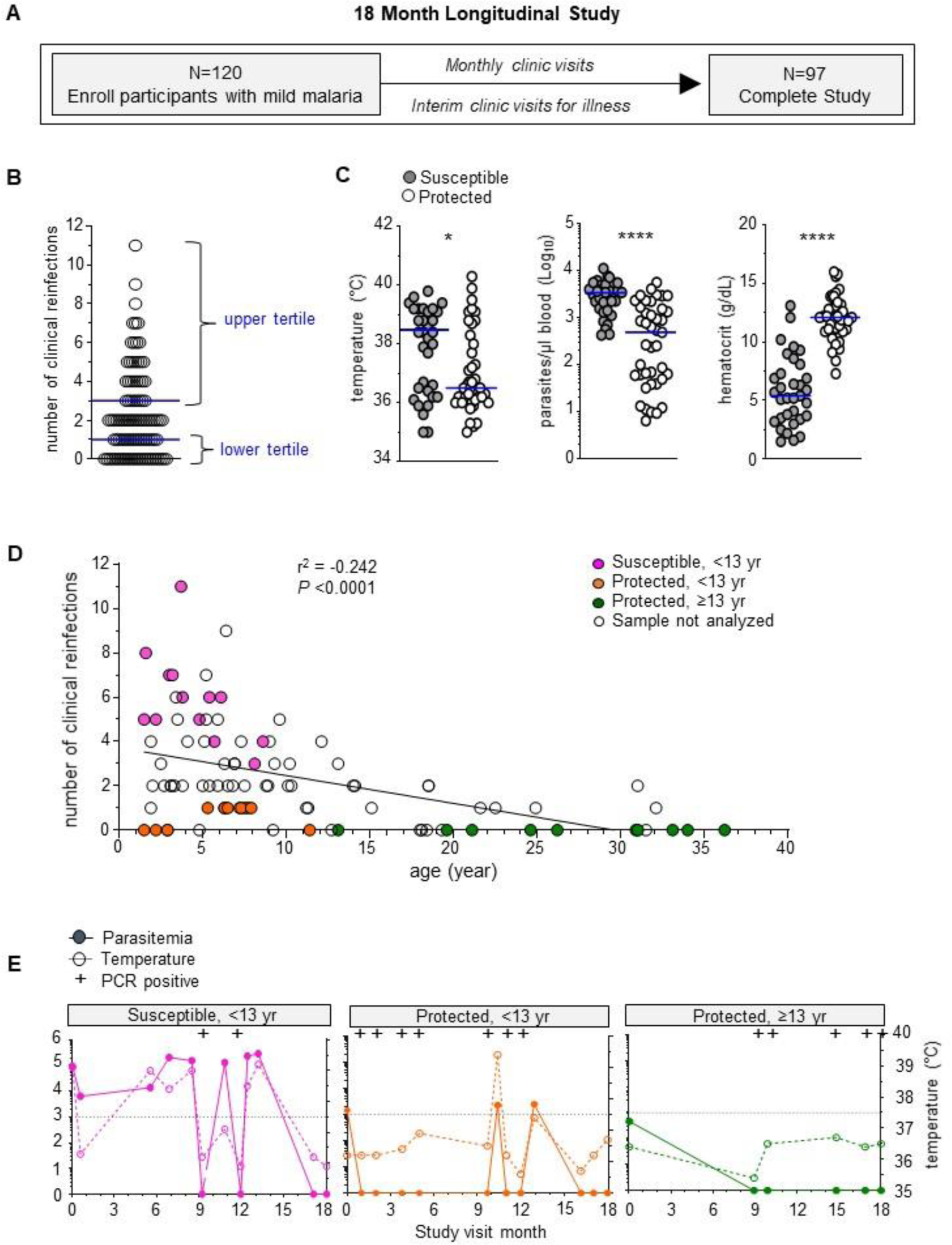
Identification of participants with varied clinical immunity against malaria in a longitudinal study of an endemic region in Malawi. (A) Overview of the 18-month longitudinal study of mild malaria participants with total enrollment number, clinical visits and retention number at the end of the study. (B) Number of clinical reinfection episodes (symptoms, >2,500 parasites/µl blood, with a prior episode at least 2 weeks apart) of each participant that completed the study. Upper tertile (67%) (≥3 clinical reinfections, susceptible, n=33) and lower tertile (33%) (≤1 clinical reinfection, protected, n=41) of participants are indicated. (C) Clinical parameters of temperature (°C), hematocrit (g/dl) and parasitemia (parasites/µl blood) of each protected (≤1 clinical reinfection, protected, n=41) and susceptible participants (≥3 clinical reinfections, susceptible, n=33) at the enrollment visit. (D) Number of total clinical reinfection episodes (y-axis, >2,500 parasites/µl blood, at least 2 weeks apart) and participants’ age at time of enrollment (x-axis). Data was stratified by age, with susceptible participants (<13 yr, ≥3 clinical reinfections), protected participants (<13 yr, ≤1 clinical reinfection) and protected participants (≥13 yr, ≤1 clinical reinfection) are indicated. Pearson correlation coefficient and statistical significance are indicated. (E) Parasitemia (left axis), temperature (°C, right axis) and *Pf* PCR status indicated for each clinical visit of one representative susceptible, one protected age-matched and one protected adult participant over the 18-month longitudinal study. Dotted line is at 37.5°C. Statistics: Student’s t-test was conducted between indicated groups, *p<0.05, **p<0.01, ***p<0.001, ****p<0.0001.

### Selection of participants who vary in clinical immunity for in depth immune analyses

The age at which clinical immunity develops within a population is based on transmission intensity and other local population level effects and thus needs to be defined in each malaria endemic region ^51, 52^. To classify Chikwawa participants as either susceptible or clinically immune we enumerated the number of recurrent clinical malaria episodes over the18-month longitudinal study. An infection was counted if the malaria genotype determined by sequence analysis was distinct from the prior infection or occurred at least two weeks after a prior infection in each participant. We then defined clinically susceptible participants as those with the greatest number of clinical reinfections (upper tertile, ≥3) and protected ones as those with the lowest number of clinical reinfections (lower tertile, ≤1) (Figure 1B). As expected, clinically susceptible participants were younger in age compared to protected individuals (Figure S1B). They had higher body temperatures, greater blood parasite loads and lower hematocrits than the protected participants at the enrollment malaria episode (Figure 1C).

To determine if the higher number of clinical infections in younger participants compared to protected individuals was due to greater exposure to malaria rather than immunity, we measured the total number of afebrile (temperature <37.5°C), low blood parasite load (< 2,500 parasites/μl) and asymptomatic reinfections and the number of microscopy negative/PCR positive reinfections over the course of the study and found no differences between the groups in either scenario (Figure S1C). This suggested that the age-related differences in the number of clinical reinfections was not simply secondary to differences in *Pf* infections, and thus likely related to immune status.

All clinically susceptible participants were less than 13 years of age. To allow comparison of immune responses among children only, we age-stratified the clinically protected participants into two groups, e.g., younger than 13 years or 13 years and older. Within these groups, we selected a set of participants for further analyses (Figure 1D). Plots of representative individuals for all three groups reporting all infections (including microscopic and submicroscopic/PCR^+^), parasitemia and temperature over 18 months of follow up are provided (Figure 1E and Figure S1D). In summary, we established a well-characterized cohort residing in a high transmission *Pf* endemic area that demonstrated variation in clinical immunity. This enabled us to conduct in depth immunophenotyping of their peripheral blood mononuclear cells (PBMCs), analysis of their plasma Ab and single cell transcriptomic to investigate host immunity associated with protection against symptomatic re-infections.

### *Pf-*specific humoral responses in malaria immune compared to clinically susceptible participants

Abs against malaria Ags play an important role against disease and are associated with clinical immunity ^31, 33, 53, 54^. As an additional benchmark for our classification of clinical immunity, we characterized the Ab response in the plasma samples from participants at enrollment during clinical malaria using a *Pf* protein array displaying ∼1,130 predicted and known *Pf* proteins ^40, 55, 56^ (Figure 2 and Figure S2). Among all subjects combined, plasma IgG Ab reactivity was found against a total of 372 *Pf* Ags, as represented in a heatmap (Figure S2A and Table S1). Overall, plasma anti-malarial IgG Abs from the older protected group reacted to a significantly higher number of *Pf* Ags (∼177 antigens) compared to the susceptible group (∼125) (Figure 2A, upper graph). However, there was no significant difference between IgG Abs reactivity to *Pf*- Ags comparing the young protected with the susceptible group. Further, plasma IgG Ab levels against individual *Pf-*Ags were significantly higher in plasma from older protected subjects compared to that from susceptible and young protected participants (Figure 2A, lower graph).

**Figure 2.**
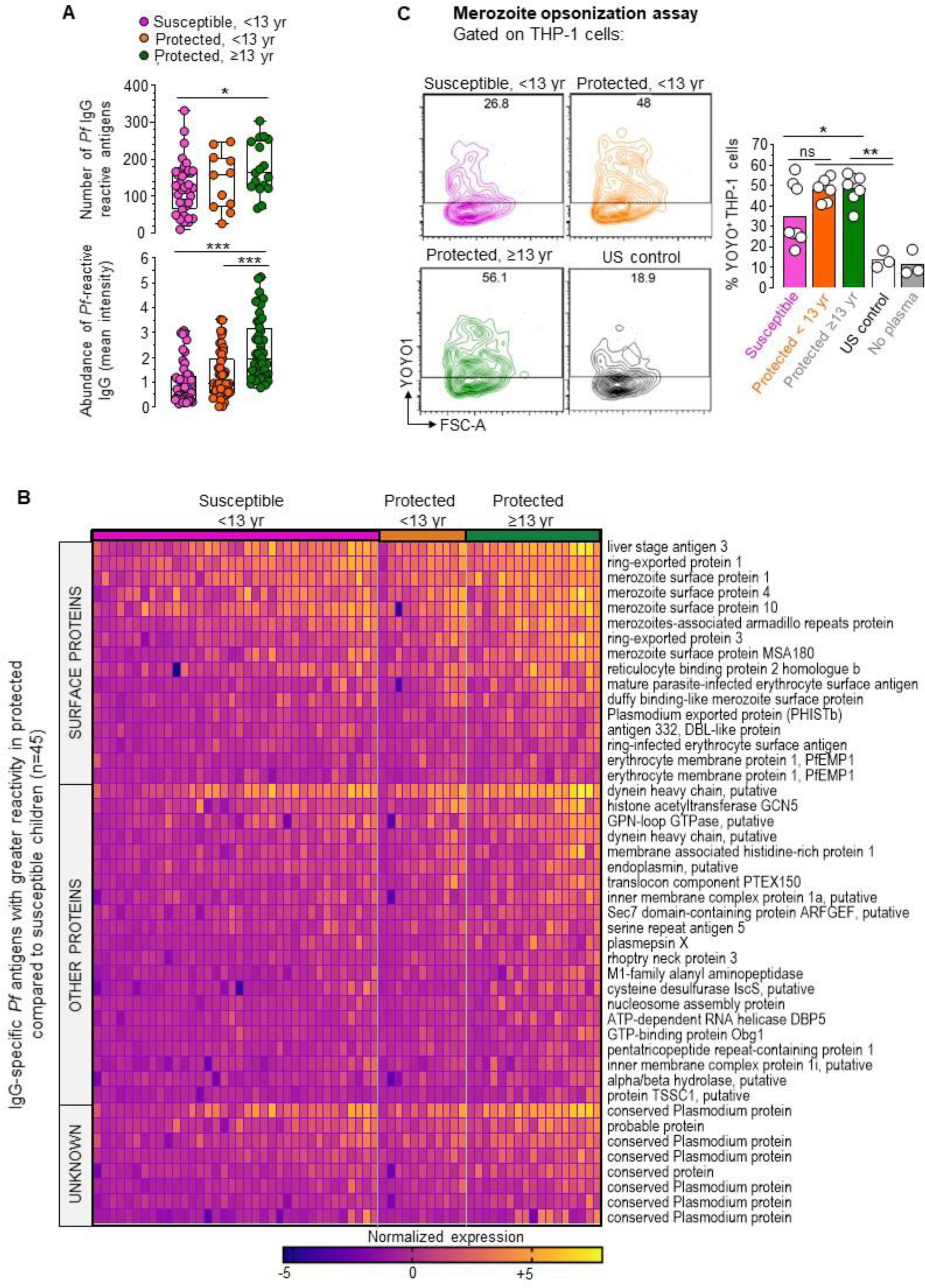
Monitoring of *Pf* specific antibody responses in participants during mild malaria illness. **(A)** Number of *Pf* antigens recognized by participants plasma IgG (top) and abundance (bottom) of *Pf* -reactive plasma IgG antibodies (mean intensity) in plasma determined on *Pf* antigen array. Median and min to max indicated on the box and whisker plots. Welch’s unpaired t-test (top) and Mann-Whitney test (bottom), was conducted between indicated groups *p<0.05, **p<0.01, ***p<0.001 **(B)** Heatmap of normalized levels of IgG antibodies in individual participants (columns) against 45 *Pf* antigens (rows) that are significantly higher in protected versus susceptible groups (Mann-Whitney test, p<0.0001). **(C)** *Pf* merozoite opsonization assay with YOYO-1 labeled *Pf* merozoites incubated with indicated participants plasma, washed and then co-cultured with THP-1 cells. FACS plots of a representative sample within indicated groups, quantifying YOYO1^+^ THP1 cells after the co-culture (left). Summary of %YOYO1^+^ THP1 cells (right) indicating % *Pf* merozoite uptake when labeled merozoites were incubated with or without indicated plasma and co-cultured with THP1 cells (n=3 to 8 plasma samples per group). Mean value indicated and Welch’s unpaired t-test were conducted between indicated groups, *p<0.05, **p<0.01, ***p<0.001.

Of note, many of the *Pf*-Ag reactive IgG Abs (n=102) were detected in all groups but they exhibited significant differences in titers (Figure S2B and Table S2). Among the 372 *Pf* reactive Ags, a significantly higher IgG reactivity was measured against 45 *Pf* Ags in the plasma from all protected compared to susceptible participants, which included 16 *Pf* surface proteins, 24 non-surface proteins and 8 unknown proteins (Figure 2B and Table S3). One of the most differentially abundant Ab reactivities was against the liver stage antigen 3 (LSA-3), which is expressed during the liver stage. Unlike other Ags, LSA-3 is highly conserved and immunization with LSA-3 induces protection in primates against heterologous *Pf* strain challenges ^57^. Plasma Abs against the merozoite stage (MSP1, MSP4, MSP10, MSA180 and Duffy binding like merozoite surface protein)^58^ were also greater in older protected individuals compared to susceptible participants (Table S3).

To determine if there were also functional differences in anti-merozoite Abs, we conducted a *Pf* merozoite opsonization assay. *Pf* 3D7 strain was grown *in vitro* under standard conditions and merozoites were isolated and labeled with the YOYO-1 fluorescent dye to detect parasite nucleic acid by flow cytometry. Fluorescently labeled merozoites were incubated with plasma for one hour, washed and co-cultured with THP-1 cells for ten minutes prior to quantification of uptake by flow cytometry (Figure 2C). Approximately 50% of THP-1 cells were YOYO-1^+^ when co-cultured with merozoites sensitized by preincubation with clinically immune participants plasma of all ages. In comparison, only 35% of THP-1 cells stained positive for YOYO-1 after co-culture with merozoites preincubated with plasma from the susceptible group. The addition of control plasma from malaria-unexposed US donors or no plasma, resulted in 10% YOYO-1^+^ staining of THP-1 cells. The *Pf* merozoite uptake was reduced in the presence of anti-Fc receptor blocking Ab, indicating that plasma incubated *Pf* merozoites uptake was Fc-receptor mediated (Figure S2C). Uptake of merozoites preincubated with plasma from clinically immune subjects was also partially blocked by cytochalasin D, showing a role for actin polymerization. The broader plasma IgG reactivity and titers against *Pf* Ags, together with the increased functional *Pf*-merozoite specific IgG responses in the protected compared to the susceptible participants, further verifies the clinical immunity classifications for the in-depth studies of their immune cells presented below.

### High dimensional CyTOF analysis of immune cell populations

To globally survey the complex cellular immune processes occurring in the peripheral blood of participants undergoing acute *Pf* malaria infection, we developed a comprehensive immunophenotyping panel of markers for mass cytometry (CyTOF) analysis of all major mononuclear myeloid and lymphoid cell lineages (Figure 3, Figure S3A and Table S4). The panel included established cell-surface and intracellular markers to discriminate effector and memory lymphocytes (CD62L, CD27, CD45RA), and several CD4^+^ T helper cell functional subsets (Foxp3, CCR4, CCR6, PD-1, CXCR5, CXCR3). Using this panel, we analyzed PBMCs from clinically susceptible participants (n=12) and the two clinically protected groups (<13 yr, n=9, and ≥ 13 yr, n=10) (Figure 1D). We processed the analysis of our high dimensional flow cytometry data using both t-Distributed Stochastic Neighbor Embedding (t-SNE) and flow Self-Organizing Map (FlowSOM) tools for unsupervised clustering and dimensionality reduction, and the visualization of discrete cell subsets and their relative proportions (Figure 3A).

**Figure 3.**
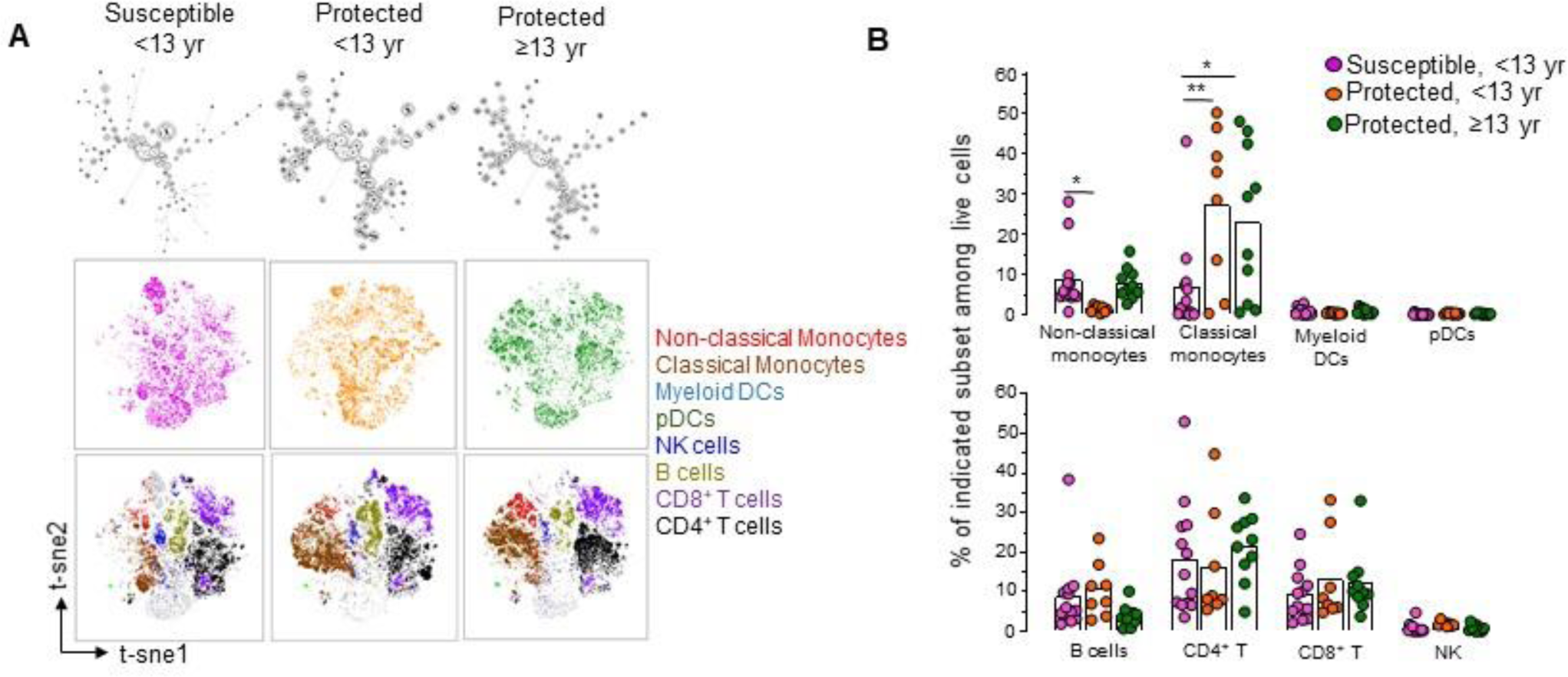
Overview of CyTOF analysis of immune cell populations in the blood of malaria infected participants. **(A)** FlowSOM (top) and t-SNE (bottom) visualization of live PBMCs of a representative sample within each indicated group based on lineage markers expressed on cells and detected by mass cytometry (CyTOF). Overlaid immune cell populations on the t-SNE plots are indicated. **(B)** Summary of innate and adaptive immune cell frequencies in PBMCs of malaria infected participants. Mean cell frequency is indicated for susceptible, <13 yr (n=13), protected, <13 yr (n=8) and protected, ≥ 13 yr (n=10). Summary of the immune cell subset frequency in participants during malaria illness among indicated groups are shown. Bar graphs represent mean cell frequency values. Statistical significance is calculated by unpaired Student’s t-test between indicated groups, *p<0.05, **p<0.01, ***p<0.001.

There was heterogeneity in the proportions of immune cell subsets in the protected versus the susceptible groups (Figure 3B and Figure S3B, C). Specifically, among innate immune cell populations, we observed significantly higher frequencies of classical monocytes (CD14^+^CD16^neg^) in both the young and older protected groups compared to the susceptible group (factor of ∼3-4), consistent with prior observations ^59^. However, non-classical, patrolling monocytes (CD14^dim^CD16^+^) were significantly reduced in the young protected group (factor of∼10) compared to the susceptible group. Similar frequencies of plasmacytoid dendritic cells (pDCs, CD123^+^) and myeloid DCs (CD11c^+^) were observed across the groups. Likewise, the proportion of total B, CD4^+^ T, CD8^+^ T and NK lymphocytes were comparable in all groups. Upon sub-analysis of these lymphocyte subsets into naïve (CD45RA^+^CD27^+^), effector (CD45RA^+^CD27^-^), and memory (CD45RA^-^CD27^+/-^) cells, there was an increased frequency of memory T cells among CD4^+^ T but not CD8^+^ T cells, in the older protected group compared to the susceptible group, which may be age-related (factor of ∼1.6 fold, Figure 4A, B) ^60^.

**Figure 4.**
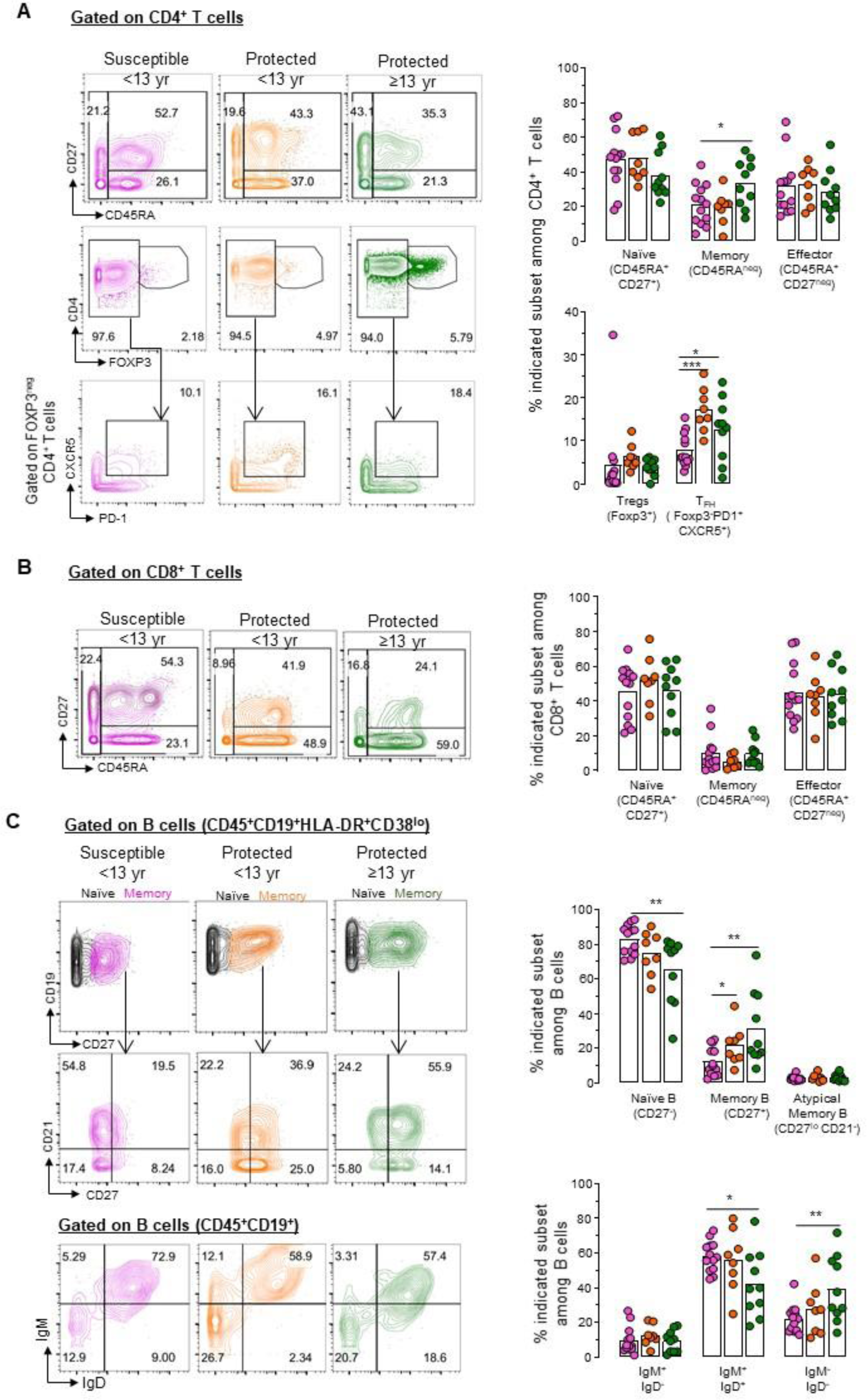
CyTOF analysis of T and B cell populations in the blood of malaria infected participants. **(A-C)** Representative CyTOF plots of one participant from each group stained with a 33 marker heavy-metal tagged antibody cocktail. Contour plots indicate the staining and gating strategy for CD4^+^, CD8^+^ T cell and B cell subsets among live PBMCs. Summary of the immune cell subset frequency in participants during malaria illness among indicated groups are shown. Bar graphs represent mean cell frequency values. Statistical significance is calculated by unpaired Student’s t-test between indicated groups, *p<0.05, **p<0.01, ***p<0.001.

Interestingly, the proportion of follicular helper CD4^+^ T (TFH) cells, but not of Foxp3^+^ regulatory T (Treg) cells, was increased in both protected groups compared to the susceptible counterpart. Consistent with this observation, we detected a significantly higher frequency of memory B cells (CD19^+^CD27^+^, factor of ∼2) among both protected groups compared to the susceptible group (Figure 4C). We also quantified increased proportions of immunoglobulin isotype class-switched B cells (IgM^-^IgD^-^, factor of ∼1.8) and reduced frequency of non-switched B cells (IgM^+^IgD^+^) by a factor of ∼1.3 when comparing these same groups. Of note, similar frequencies of atypical memory B cells (CD27^-^CD21^-^), which were reported to be expanded in malaria-infected patients, were measured across all groups ^11, 61, 62^. In summary, both clinically protected groups undergoing an episode of *Pf* infection exhibited a significantly expanded population of classical monocytes, CD4^+^ TFH and class-switched B cells during an episode of mild malaria compared to the clinically susceptible group.

### Single-cell transcriptomic analysis of circulating memory CD4^+^ T cells in protected participants

Since CD4^+^ T cells are essential to the control of *Pf* blood stage infections and the induction of long-term protective parasite-specific humoral responses ^44^, we next sought to further characterize the memory CD4^+^ T cells present in the clinically protected group during an episode of mild malaria, using single cell transcriptomic and T cell receptor (TCR) sequencing. Using high speed Fluorescence activated cell sorting with gating on live cells, memory CD4^+^ T cells (CD45RO^+^ CD27^+^ or CD27^-^) and naïve CD4^+^ T cells (CD45RO^-^CD27^+^) were purified from the PBMCs of three participants in the clinically protected group (Figure S4A). Whole transcriptome single-cell RNA-sequencing was performed on the sorted populations using the 10x Genomics platform. We integrated the single cell transcript expression data from all three participants and conducted dimensional reduction analysis to visualize transcriptionally distinct cell subsets in a UMAP plot based on transcriptional similarities between cells (Figure 5A). This identified eight transcriptionally distinct memory CD4^+^ T cell subsets, found across the three participants analyzed. The distribution of subsets within individual participants was similar (Figure S4B). As a control, we examined naïve CD4^+^ T cells which exhibited a very different transcriptional profile and clustered separately from memory CD4^+^ T cells across all samples, as visualized on the UMAP plots (Figure S4C, D). None of clusters found in the memory CD4^+^ T cell population overlapped with that of the naïve (Table S5). This differential clustering of naïve versus memory CD4^+^ T cells was also not driven by differences in participant samples, since each individual sample had a similar subset distribution across all clusters for both the naïve and memory populations (Figure S4B, E).

**Figure 5.**
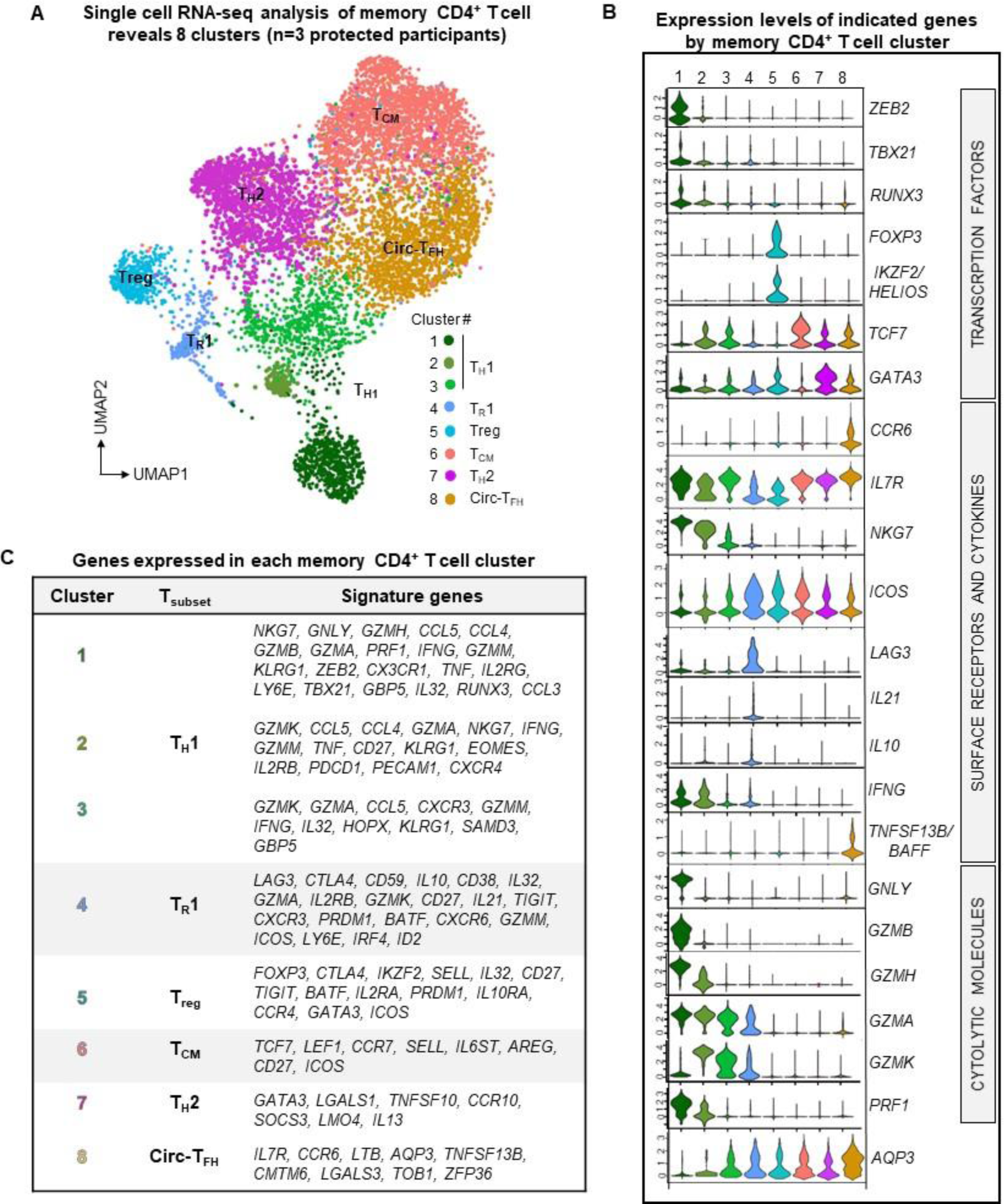
Single-cell transcriptomic analysis of memory CD4^+^ T cells from protected participants during malaria illness. **(A)** UMAP visualization of integrated single-cell RNA-seq results on memory CD4^+^ T cells isolated from 3 protected participants during malaria infection. Clusters of memory CD4^+^ T cells with distinct transcriptional profiles are colored as indicated in the legend. **(B)** Normalized expression levels of indicated genes within each distinct memory CD4^+^ T cell cluster defined in (A). **(C)** Table of top signature genes expressed in each distinct memory CD4^+^ T cell cluster defined in (A).

Among clusters of memory CD4^+^ T cells, clusters 1, 2 and 3, showed a robust TH1 expression profile (Figure 5B, C). Cluster 1 compared to 2 and 3, exhibited significantly higher levels of transcripts coding for T-bet (*TBX21*), the master Th1 transcriptional regulator, and for Zeb-2, a key transcription factor (TF) that cooperates with T-bet to drive terminal effector CD8^+^ T cell differentiation ^63, 64^. Moreover, cluster 1 expressed higher levels of transcripts coding for the TF Runx3, which epigenetically programs and enables the differentiation of long-lived cytolytic memory T cells ^65, 66^. We also noted upregulation of genes in cluster 1 encoding multiple effector molecules including granzymes (*GZMB*, *GZMA*, *GZMK*, *GZMH*), granulysin (*GNLY*), perforin (*PRF1*), IFNγ, as well as *NKG7* which encodes a membrane protein that regulates target cell killing ^67^. Clusters 2 and 3 selectively expressed transcripts coding for granzyme K (*GZMK*) but not B (*GZMB*), compared to cluster 1. Other notable differences between cluster 2 and 3 were the abundance of granzyme H, IFNγ and perforin-encoding transcripts (Figure 5B). While these 3 clusters have a clear TH1 cell signature, they may represent distinct stages or paths of TH1 cell differentiation.

In addition, the single cell transcriptomic analysis identified subsets of regulatory T (Treg) cells. These included a population that were most likely TR1 cells (cluster 4), that expressed transcripts of inhibitory receptors (LAG-3, CTLA-4, ICOS, TIGIT) and the cytokines IL-10 and IL-21, while also encoding mRNA of TFs involved in terminal effector cell differentiation (*PRDM1, ID2*) ^47, 48, 68^. It also included Treg cells that expressed increased transcript levels of the TFs Foxp3 and IKZF2 (Helios), and inhibitory receptors CTLA-4, ICOS, and TIGIT (cluster 5). Among the largest subsets revealed by this analysis, we found gene expression signatures of central memory cells (TCM, cluster 6) and TH2 effector cells (cluster 7) with high levels of transcripts coding for their respective master transcriptional regulators TCF7 and GATA-3. Lastly, we report an abundant subset of memory CD4^+^ T cells (cluster 8) that contained high levels of transcripts coding for the TF TCF7, the chemokine receptor CCR6, the cytokine BAFF and the IL7 receptor, consistent with a signature of circulating TFH cells (Circ- TFH) ^10, 68, 69^.

### Expanded memory CD4^+^ T cell clones expressing shared TCR clonotypes have a cytolytic transcriptional signature

Concomitantly with the single cell transcriptomic analysis, we sequenced the TCRα and TCRβ variable chains expressed in each memory CD4^+^ T cell sorted from *Pf*-infected protected participant PBMCs (Figure 6 and Figure S5). With this approach, we could determine if individual memory CD4^+^ T cells expressed exactly the same pair of TCRα and β chain sequences and thus shared the same TCR clonotypes (Figure 6A and Figure S5A). Between ∼0.3% to 4.3% of memory CD4^+^ T cells bear identical TCR clonotypes in each participant, suggesting that each of these expanded T cell clones are the progeny of the same original clone expanded in response to a specific Ag. Since the TCR sequencing analysis was conducted on the memory CD4^+^ T cells for which we ran our transcriptomic analysis, we could link each expanded T cell clone to its expression signature and the memory CD4^+^ T cell clusters defined in Figure 5A (Figure 6B). The large majority (∼95%) of expanded T cell clones sharing a same TCR clonotype across the three participants belonged to cluster 1 -which expressed transcripts encoding a robust cytolytic TH1 effector program (Figure 6C). All expanded clones expressed high levels of transcripts coding for master transcriptional regulators governing effector T cell signature and fates, namely Zeb2 (∼60%), Runx3 (∼35%) and T-bet (∼20%) (Figure S5B). These cells also expressed transcripts coding for the cytolytic effector proteins NKG7 (∼100%), granulysin (*GLNY*, ∼93%), granzymes (*GZMH, GZMA, ∼*94%), granzyme B (*GZMB,* ∼70%) and perforin (*PRF1,* ∼80%). Furthermore, clusters 2 and 3 (TH1, GMZK^+^, GZMB^-^GNLY^-^), and cluster 4 (TR1) respectively, accounted for ∼2% of the expanded clones (Figure 6C).

**Figure 6.**
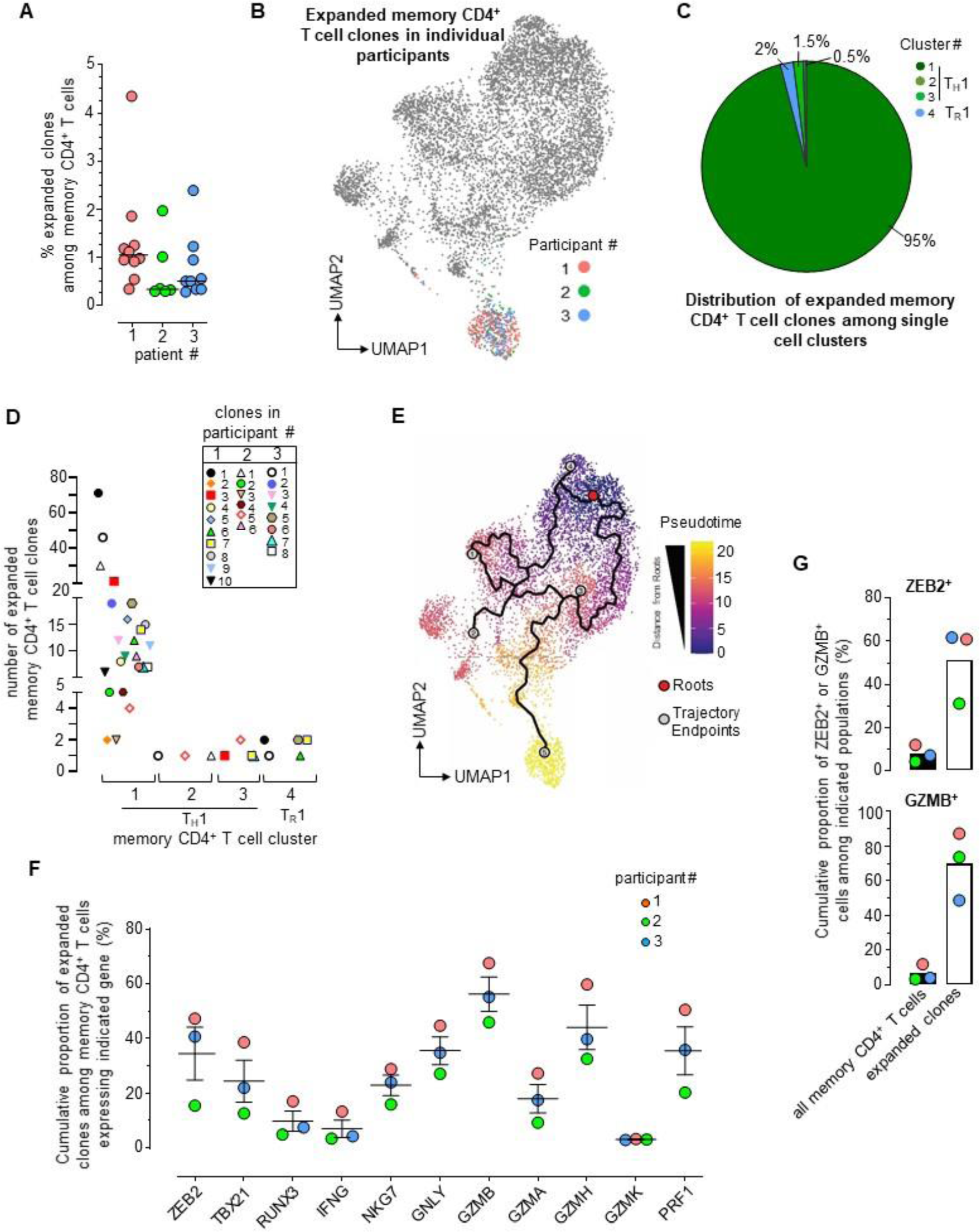
Expanded TCR clones among memory CD4^+^ T cells in malaria protected patients exhibit a cytolytic gene expression signature. **(A)** Frequency of each expanded memory CD4^+^ T cell clone expressing an identical TCR clonotype among all memory CD4^+^ T cells in indicated participant, as defined by single-cell TCR-seq. **(B)** Expanded memory CD4^+^ T cell clones (at least 10 cells sequenced with the exact same TCRα and TCRβ chains) in each of the 3 participants, highlighted on the concatenated UMAP visualization shown in Figure 5A. **(C)** Pie- chart of expanded memory CD4^+^ T cell clone proportions across distinct transcriptional clusters defined in figure 5A and across the 3 protected participants. **(D)** Distribution of individual clones with identical TCR clonotype across memory CD4^+^ T cell cluster 1, 2, 3 and 4 (as defined in Figure 5) and across the 3 protected participants. **(E)** Pseudotime analysis of the memory CD4^+^ T cell subsets of Figure 5A. **(F)** Cumulative proportion of expanded memory CD4^+^ T cell clones expressing indicated gene-encoding transcripts across the 3 participants. **(G)** Proportion of cells that express *ZEB2* or *GZMB* transcripts among all memory CD4^+^ T cells or only among expanded T cell clones.

To further understand if expanded T cells expressing the same TCR clonotypes could be found in the multiple memory CD4^+^ T cell clusters defined in Figure 5A, we next tracked the fate of each of them across the various clusters (Figure 6D). Several clones sharing the same TCR clonotype, belonged to various clusters (1 and 2 ; 1 and 3 ; 1 and 4 ; 1, 2 and 3 ; 1, 2 and 4 ; 1, 3 and 4), suggesting that an expanded clone can have multiple TH fates. To assess individual clone fates in the context of T cell differentiation, we defined single cell trajectories using Monocles ^71^. Monocles models single cell gene expression data as a function of pseudotime to place a cell along the differentiation process (Figure 6E). Pseudotime is an arbitrary measure reflecting how far an individual cell is in the differentiation process with respect to a predefined cell of origin (roots). Here we used TCM as the root (cluster 6, Figure 5A) since these cells are well-reported to maintain the highest differentiation potential among memory cells. Together with their individual distribution among the relevant clusters (1 to 4, Figure 6E), this analysis suggested that the most highly differentiated memory CD4^+^ T cell subset (TH1, cluster 1), where expanded clones accumulated the most, likely derived from less differentiated states that included TR1, TH1 cluster 2 or TH1 cluster 3. Taken together these results show that the great majority of expanded memory CD4^+^ T cell clones sharing identical TCR clonotypes in clinically protected individuals during mild *Pf* malaria infection, were terminally differentiated and expressed a robust cytolytic effector program.

To identify potentially unique biomarkers of the expanded clones, we defined the clonotype transcriptional signatures and compared their proportion among all memory CD4^+^ T cell transcripts (Figure 6F and Figure S5C). Among the transcripts encoding for transcriptional regulators, *ZEB2* was the most highly expressed in each of the expanded clones, and ∼35% of the *ZEB2*-expressing memory CD4^+^ T cells were expanded clones. In addition, among the transcripts coding for cytolytic effector molecules, *GZMB* expressing memory CD4^+^ T cells consisted of the largest proportion of expanded clones among memory CD4^+^ T cells (∼60%). Both *ZEB2*- and *GZMB*-expressing memory CD4^+^ T cells were significantly enriched in expanded clones compared to that of memory CD4^+^ T cells (factor of 7-12, Figure 6G). These results collectively suggested that a substantial proportion of memory CD4^+^ T cells that express *ZEB2* or *GZMB* are clonally expanded and presumably respond to *Pf* antigen, raising the possibility that expression of Zeb2 and granzyme B may be useful as surrogate markers to identify malaria-specific memory CD4^+^ T cells that are clonally expanded.

### Expression of multiple chemokine and inhibitory receptors by Zeb2^+^ memory CD4^+^ T cells

Using high dimensional spectral flow cytometry, we confirmed that memory CD4^+^ T cells expressed Zeb2 and granzyme B and characterized them further using multiple cell surface markers based on our single cell analysis (Figure S6A). Tracking of Zeb2^+^ cells among memory CD4^+^ T cells in the blood of susceptible and young protected groups revealed that these cells are significantly expanded at the time of acute infection compared to 30 days later at convalescence (Figure 7A). While Zeb2^+^ cells were detected in older clinically protected participants at equivalent frequencies as in other convalescent participants, we did not detect an expansion of the Zeb2^+^ population during illness in this older protected group, which was in contrast to our observation in the younger groups. Unlike the Zeb2^+^ memory CD4^+^ T cells, we did not detect any expansion of granzyme B^+^ cells during infection compared to convalescence, suggesting that the expression of granzyme B may not sufficiently track with the expanded clones (Figure S6B).

**Figure 7.**
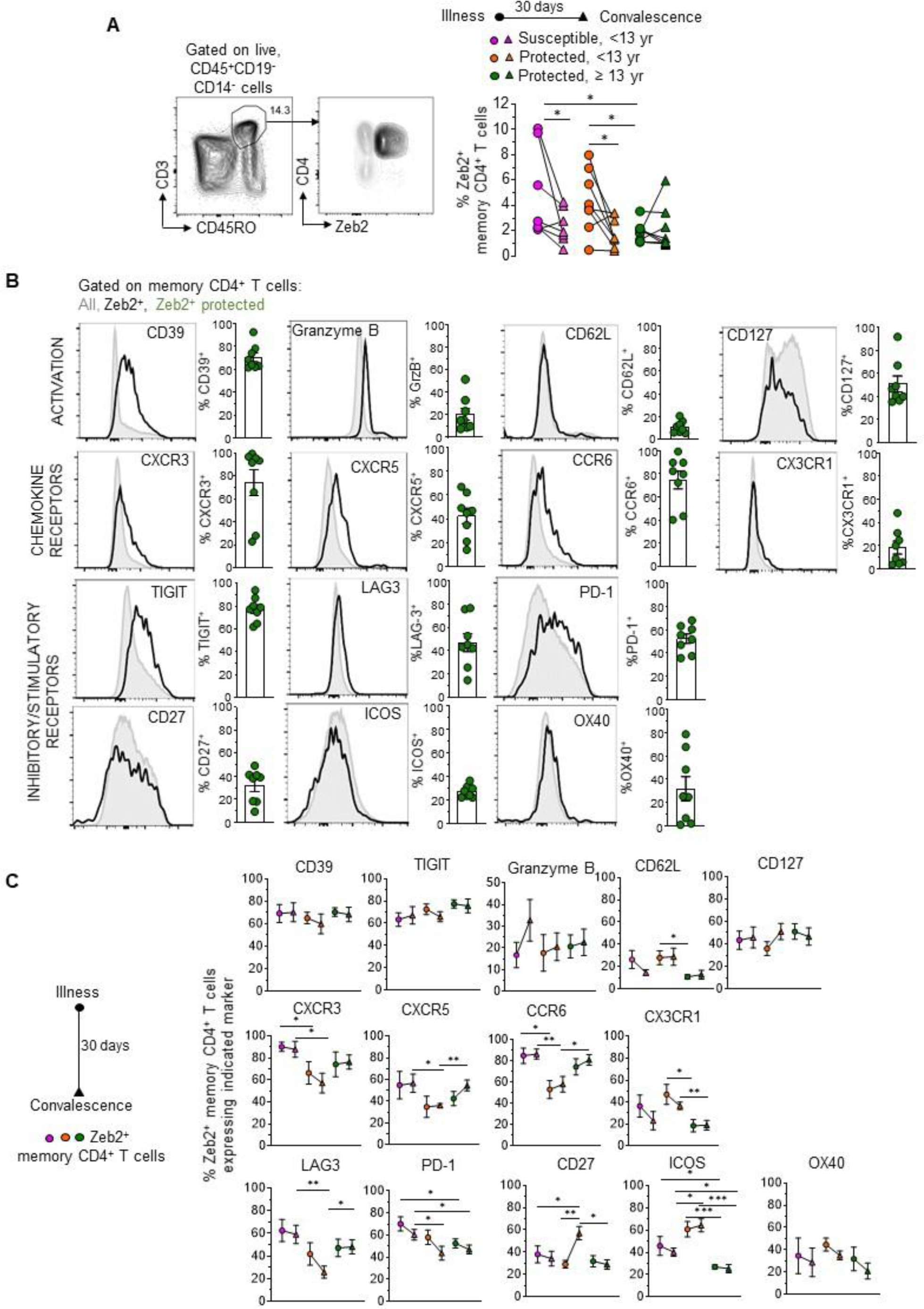
Characterization of Zeb2^+^ memory CD4^+^ T cells during and after a malaria episode. **(A)** Gating strategy of Zeb2^+^ memory CD4^+^ T cells using spectral flow-cytometry on a representative protected malaria participants during illness. Frequency of ZEB2^+^ memory CD4^+^ T cells in susceptible, <13 yr (n=7, pink), protected, <13 yr (n=8, orange) or protected, ≥13 yr (n=8, green) malaria participants during illness (circle symbol) and at day 30 post illness (convalescence, triangle symbol). **(B)** Representative FACS histograms of the expression levels of indicated markers on Zeb2^+^ (black line) or all (grey line) memory CD4^+^ T cells in a representative protected participant. Bar graphs average the proportion of memory cells expressing each marker in protected participants, and each symbol (green dots) represents 1 participant. **(C)** Frequency of indicated markers on Zeb2^+^ (filled symbol) memory CD4^+^ T cells in susceptible (pink, n= 8), young protected (orange, n=7) or old protected (green, n=8) malaria participants during illness and at day 30 post illness. Mean and SEM values for each group are shown. Statistics is a Wilcoxon paired t-test for comparisons within each group and a Student’s t- test for comparisons across groups as indicated, *p<0.05, **p<0.01, ***p<0.001.

Further analysis of the phenotype of Zeb2^+^ memory CD4^+^ T cells during infection in older protected participants revealed that these cells were effector memory cells (CD62L^low^CD127^low^), contained high amounts of granzyme B and expressed high levels of CD39, a surrogate marker upregulated on recently activated antigen-specific cells (Figure 7B and Figure S6C).

Zeb2^+^memory CD4^+^ T cells also expressed measurable levels of cell-surface chemokine receptors (CXCR3, CXCR5, CCR6) further suggesting that they may better respond to multiple chemotactic inflammatory cues. We also noted that they generally expressed low levels of cell-surface costimulatory molecules (ICOS, CD27, OX40) while inhibitory receptors (TIGIT, LAG- 3, PD-1) were upregulated, suggesting they may be subjected to various mechanisms of immune regulation.

Among these markers, expression of CD39 and TIGIT was constant between groups during illness and convalescence. However, significant differences in the phenotype of Zeb2^+^ memory CD4^+^ T cells were also observed between groups during illness and convalescence. We report significant differences in older protected compared to susceptible participants for expression of ICOS and PD-1, which are markers also associated with TFH cells. We also found a higher proportion of CX3CR1^+^ cells and a lower proportion of CXCR3^+^, CXCR5^+^ and CCR6^+^ cells in young protected compared to susceptible participants. Likewise, we also measured clearly distinct proportions of costimulatory (ICOS, OX40, CD27) and inhibitory (LAG-3, PD-1) receptor-expressing Zeb2^+^ memory CD4^+^ T cells. Since the phenotype of these cells in both protected groups differed significantly, this also suggested that young protected participants may be relying on immune mechanisms of protection that are distinct than those present in older protected individuals ^72^ . Collectively, these data suggest that Zeb2^+^ memory CD4^+^ T cells are expanded during an episode of malaria and express chemotactic and inhibitory receptors, which may represent key functional features associated with natural anti-malarial protective CD4^+^ T cell responses in human malaria.

## DISCUSSION

In this work, we conducted a longitudinal study to monitor study participants living in a malaria-endemic areas with high, year-round transmission, where residents develop naturally acquired clinical immunity against severe disease and clinical illness during malaria infection. We postulated that a comparison between individuals who have or have not yet developed clinical immunity would reveal new functional immune cell subsets associated with natural protection that may ultimately inform the design of an efficacious malaria vaccine. We present a detailed phenotypical classification of study participants who vary in clinical immunity to provide a robust foundation for our immune analyses. During an episode of mild malaria, we found that protected individuals were less likely to be febrile, had lower blood parasite loads, and higher hematocrit. These individuals exhibited greater parasite-specific Ab responses (breadth, titers, opsonization) and higher proportion of classical monocytes, switched IgG^+^ B cells and CD4^+^ TFH cells. Importantly, we report the identification of a subset of highly differentiated memory CD4^+^ T cells that are clonally expanded during clinical malaria, express the Zeb2 transcriptional regulator and exhibit a robust cytolytic effector gene signature. These cells also express high levels of cell-surface CD39 and TIGIT, and multiple chemotactic and inhibitory receptors.

The presence of cytolytic CD4^+^ T cells in malaria-infected patients and mouse models of malaria was first reported several decades ago ^41, 73^. Building on the hypothesis that CD8^+^ T-cell dependent cytolytic activity could play a major role in anti-sporozoite immunity, these original studies isolated MHC class II-restricted cytolytic CD4^+^ T cell clones that recognized a *Pf* circumsporozoite (CS) protein-derived epitope (human) and a *P. berghei* epitope common to both the liver and blood stage infection of this rodent parasite. In both studies, these clones could expand, produce IFNγ and effectively lyse parasite Ag-pulsed target cells. The mouse model further established that transfer of these cells protected mice against infection with a live sporozoite challenge. In these reports, the CD4^+^ T cell cytolytic clones were isolated from the animal model of malaria or human subjects that received irradiated sporozoite immunizations.

The presence of parasite (CS)-specific CD4^+^ T cells with cytolytic activity was more recently confirmed in individuals inoculated with irradiated or live sporozoite immunizations ^42, 43, 74^. To our knowledge, however, no studies have reported the presence of such cells in individuals who have developed natural immunity against *Pf*. This finding is particularly interesting as sporozoite immunization induces potent protection against homologous *Pf* challenge infections and is viewed as a promising vaccine strategy that could induce high levels of immunity ^26, 28, 29, 75, 76^.

While natural immunity is not sterilizing, it still effectively lessens the symptoms of clinical malaria episode and prevents severe disease, and it is long-lived. The Zeb2^+^ cytolytic memory CD4^+^ T cells we described are clonally expanded in the blood of clinically immune individuals. We also confirmed their overall expansion among memory CD4^+^ T cells in the blood of susceptible and young protected groups undergoing an episode of acute malaria compared to convalescence. This was not observed in the older clinically immune group and could be the result of differences in the kinetics and localization of these cells. The immune response in vaccinated or immune individuals is known to be faster and more potent upon infection, therefore enabling more rapid containment of microbial pathogens which could involve the Zeb2^+^ cytolytic memory CD4^+^ T cells.

Many studies have reported the existence of cytolytic CD4^+^ T cells, in large part during human viral infections ^77^. These cells are prominent in viral infections that target HLA class II-expressing cells such as CMV, EBV, Dengue and HIV, and which include antigen-presenting cells (DCs, B cells) and CD4^+^ T cells ^78–80^. In influenza and hepatitis C viral infections too, lung epithelial cells and hepatocytes upregulate cell-surface expression of HLA class II molecules, becoming targets for cytolytic CD4^+^ T cells ^81^. During viral infections (SIV, HIV, Influenza), these cytolytic CD4^+^ T cells are proposed to contribute to the control of viral replication through multiple effector mechanisms including direct cytolysis of infected cells. In malaria, T cell-dependent cytolysis has mostly been invoked as a mechanism to protect against liver stage infection, through CD8^+^ T cell-mediating the rapid killing of MHC class I-expressing hepatocytes infected by *Pf* sporozoites ^82, 83^. Yet, the liver stage of infection is time limited (up to a week), with a very low proportion of infected hepatocytes since mosquitoes only deliver a few hundred sporozoites into the host dermis, making it unlikely that memory CD8^+^ T cells developed in the course of natural *Pf* infection could confer significant levels of protection. It has been shown, however, that irradiated sporozoite immunizations ^26, 76^, and *Plasmodium* strains genetically engineered to arrest at late pre-erythrocytic stages ^32^, are able to promote high levels of parasite-specific immunity, likely through the production of high numbers of liver-resident parasite-specific memory CD8^+^ T cells ^84^. Thus, upon repeated exposure to *Pf* infection, it is conceivable that sufficiently high numbers of *Pf*-specific memory CD4^+^ T cells with cytolytic features may accumulate systemically and perhaps reside in the liver of infected individuals, to help rapidly control sporozoite liver re-infections prior to the development of the blood stage infection. The memory CD4^+^ T cells we describe in this study express high levels of transcripts encoding for granulysin and multiple other effector molecules (granzymes, perforin, NKG7, IFNγ), as well as chemotactic receptors (CXCR3, CCR6) that may help their rapid migration or residency next to sites of injury where sporozoites first enter. *Plasmodium* sporozoites were indeed shown to glide along the liver sinusoids until they pause and traverse Kupffer cells or liver sinusoid endothelial cells to reach hepatocytes ^83^. Kupffer cells express high levels of MHC class II and it is possible that upon recognition of parasite Ags, Zeb2^+^ memory CD4^+^ T cells arrest and release granulysin which could directly lyse paused sporozoites. Several studies suggested that CD8^+^ T cells can use such mechanism to lyse *Mycobacterium* bacteria and *P. vivax* parasites ^85, 86^. Thus, it is tempting to speculate that these cells may represent important contributors of clinical immunity against *Pf* malaria *in vivo*. Further investigations will be needed to define the antigenic specificity of these cells and the parasite Ags they recognize. Such studies will need to overcome the present lack of reagents to track Ag-specific CD4^+^ T cells in malaria, due to the tremendous HLA class II diversity, parasite Ag variability and difficulty in characterizing strong *Pf*-reactive epitopes across human populations.

More than 95% of the expanded clones sharing identical TCR clonotypes in protected participants during a malaria illness, expressed a robust TH1 cytolytic effector program including transcripts encoding multiple lytic effector proteins and the TF Zeb2, T-bet and Runx3 (cluster 1). The transcription factor (TF) Zeb2 was shown to drive the onset of terminally differentiated KLRG1^hi^ effector CD8^+^ T cells, likely under the transcriptional control of T-bet ^63, 64^. Expression of T-bet and Runx3 can turn off ThPOK and transcriptionally re-program mature CD4^+^ TH cells, the key TH cell master TF that suppresses the cytolytic program in CD4^+^ T cells ^87^. These prior findings are consistent with the transcriptional signature of expanded clones in cluster 1. Another ∼1.5% and ∼0.5% of the expanded clones belonged to TH1 cluster 2 and 3, respectively, which differentially express transcripts encoding granzyme K and Zeb2. Pseudotime analysis suggests that the expanded memory CD4^+^ T cell clones in clusters 2 and 3 represent intermediate, less differentiated stages, before they develop into robust cytolytic Zeb2^+^ memory cells. In particular, cluster 2 expanded clones express Eomes and IL-15 receptor beta chain transcripts, which regulates the expression of granzyme B in T cells. From the expanded clones, we also found that 2% had a gene expression signature consistent with that of TR1 cells (cluster 4). Pseudotime analysis suggests that cluster 4 cells are less differentiated than clusters 1-3, and are on a distinct developmental trajectory. In the mouse model of *P. berghei* malaria, highly proliferative CD4^+^ TH1 intermediates exhibit either a TH1 or a TFH fate ^68^, and some TH1 cells also initiate the expression of genes associated with TR1 cells, consistent with these findings. While we did not detect any expanded clones sharing identical TCR clonotypes in the circulating memory TFH cells, it is possible that these cells are retained in lymphoid organs and thus were not detected in the peripheral blood samples. The CyTOF analysis revealed a higher proportion of circulating memory CD4^+^ TFH cells in protected groups, which may also reflect the overall increase of this compartment in protected individuals. Further investigations are clearly needed to establish whether the Zeb2^+^ cytolytic memory CD4^+^ T cells expand and preferentially migrate to the peripheral blood while TFH counterparts remain in lymphoid organs and thus were not identified in our analysis. The significantly higher proportion of classical monocytes (CD14^+^CD16^-^) in protected participants compared to susceptible participants that we and others ^59, 68^ reported may influence the differentiation of the Zeb2^+^ memory cells, as proposed in the mouse study ^68^.

In summary, this work includes a carefully defined classification of clinical immunity, and the application of advanced analytic methods including mass cytometry and single cell analysis to provide novel insights into protective immune responses during natural *Pf* infection ^88^. We provide one of the most comprehensive analyses of immune responses during mild malaria infection and expand upon prior focused studies to identify memory CD4^+^ T cell subsets associated with clinical immunity ^11, 89–94^. We report the first single cell analysis of CD4^+^ T cell responses during human natural malaria infections for a granular analysis that defines subsets, characterize their potential function and track expanded clones sharing identical TCR clonotypes. This approach has recently been conducted in the mouse model of malaria ^68, 69^ and expands upon prior whole blood transcriptional analysis to provide novel insights into key adaptive cellular subsets and potential functions during the human infection ^88^. Thus, our study has identified humoral and cell mediated signatures associated with clinical immunity, and the identification of a clonally expanded subset of Zeb2^+^ memory CD4^+^ T cells with a robust cytolytic phenotype. Signatures of immunity may vary by malaria endemicity, host and parasite factors and malaria control interventions. Similar analyses in other regions could provide further insights into the importance of these cells in malaria immunity ^95–97^.

## MATERIALS AND METHODS

### Study Design

To identify immune correlates of clinical immunity in a high transmission malaria endemic region we recruited study participants who presented with mild malaria into a prospective 18- month longitudinal study to enumerate malaria reinfections over time at the Mfera Health Center in Chikwawa District under the auspices of the Malawi International Center of Excellence for Malaria Research ^98^. The Chikwawa District of Malawi is a wet, low-lying rural district in the Shire Valley with high, year-round malaria transmission ^99^.

Enrollment criteria required an episode of mild malaria which was defined by the WHO as presenting symptoms compatible with mild malaria and without signs of severe malaria. Participants had positive rapid diagnostic test for malaria (SD BIOLINE, Malaria Ag P. f., Abbott) at the field site, which was later confirmed positive by microscopy. Patients with chronic medical conditions including a history of HIV infection were excluded from enrollment. To capture variation in naturally acquired clinical immunity, 40 participants in each of 3 age groups (1-5, 6-12 and 13-50 years) were enrolled. After the enrollment visit, study participants underwent monthly clinic study visits and interim visits during illness to evaluate symptoms of malaria, record temperature, collect a blood smear and a blood drop onto a FTA classic cards (Whatman, NJ) for PCR detection of *Pf* ^98^. Recurrent clinical malaria was defined as an episode with clinical symptoms consistent with malaria and a positive malaria blood smear (>2,500 parasites/μl) ^100^. Parasitemia was determined by microscopy and was recorded as the geographic mean value of two independent readers. Whole blood samples were collected in at in ethylenediaminetetraacetic acid (EDTA) coated tubes at enrollment, during each subsequent episodes of clinical malaria and day 30 post infection. Within 6 hours of collection, the blood sample was centrifuged to collect plasma aliquots and isolate PBMCs using ACK lysis buffer (Crystalgen) with standard methods ^101^. PBMC aliquots were stored in 5% DMSO and 95% FBS (Fisher Scientific) in liquid nitrogen. PBMC and plasma samples were shipped to Albert Einstein College of Medicine in New York, in liquid nitrogen gas and stored in liquid nitrogen for subsequent immunophenotyping and protein array analysis. Filter papers were collected during each episode of clinical malaria and routinely at each monthly visit for parasite genotyping ^98^. An infection was counted if the malaria genotype by sequence analysis was distinct from the prior infection or occurred at least two weeks after a prior infection in each participant (including episodes of non-clinical malaria and submicroscopic infections). Participants who were diagnosed with clinical malaria were treated with 3 days of artemether/lumefantrine per Malawi Ministry of Health National Treatment Protocol. Participants were considered lost to follow-up if they did not present for > 3 consecutive monthly study clinic appointments.

Informed consent was obtained prior to study enrollment. Institutional Review Board approvals were obtained from the Albert Einstein College of Medicine, Michigan State University, the University of Maryland, and the University of Malawi College of Medicine Research and Ethics Committee.

### Detection of sickle cell polymorphism

Sickle cell polymorphism was an exclusion criteria for further immune analysis. To determine sickle cell status, blood was spotted onto FTA classic cards (Whatman, NJ) from the enrollment whole blood sample in EDTA tubes. Whole DNA was extracted using QIAamp DNA mini kit (Qiagen, Hilden Germany) following the manufacturer’s instructions. Following whole DNA extraction, the sickle cell allele status was determined by restriction fragment length polymorphism (RFLP) on PCR amplified DNA and gel electrophoresis following previously published protocols ^102^. Genomic DNA (gDNA) from a patient homozygous for the sickle cell trait was used as a positive control.

### Immunophenotyping by cytometry by time of flight (CyTOF) and spectral flow-cytometry

Frozen vials of PBMCs were placed in a 37°C water bath and gently agitated until ∼90% was thawed. The vials were transferred to a tissue culture hood and 1mL of sterile warmed media (RPMI/20%FBS) was added to the PBMCs for 5 minutes before transferring drop-wise to a 10ml volume of warmed media (RPMI/20%FBS) containing 200U/ml of DNase I (Roche, Switzerland). After 10 minutes at room temperature, the PBMCs were centrifuged at 300g, at room temperature for 5 minutes. The pellets were washed twice with room temperature media (RPMI/20%FBS) before transferring to 96 well plates for staining with antibodies for CyTOF or spectral flow cytometry.

#### For mass cytometry

PBMC samples were labeled with cisplatin, stained with heavy-metal conjugated Abs listed in Table S5, and barcoded following the manufacturer’s protocol (Fluidigm, US) with minor modifications. Abs not commercially available as conjugated with specific heavy metal ions were conjugated using the ready-to-label antibody format and Maxpar Antibody Labeling kit (Fluidigm, US). Briefly, cells were stained with extracellular Abs in Maxpar staining buffer for 30 minutes on ice followed by addition of cisplatin (5μM in PBS) for 2 minutes at room temperature. Cells were washed twice in Maxpar staining buffer followed by fixation and permeabilization using the eBioscience FoxP3/transcription factor staining kit (Thermofisher Scientific) for intracellular staining and barcode labeling. Lastly, barcoded samples were combined for intercalation using Cytofix/Cytoperm buffer (BD) and 2% PFA/Ir solution for 30 minutes at room temperature. This was followed by a wash and resuspension of cells in MaxPar cell staining buffer with 125nM of Ir and storage at 4°C before running on the Helios CyTOF instrument (Fluidigm) at the Icahn School of Medicine Mount Sinai, Human Monitoring Core (New York, NY). The data was analyzed on FlowJo LLC software (v10.7.1, BD, Ashland OR).

#### For spectral flow cytometry

participant’s PBMCs were first stained with Live/dead Fixable Aqua (Thermofisher Scientific) for 30 minutes at room temperature. Cells were washed and incubated with human Fc block (BD) for 15 minutes on ice, followed by staining for extracellular markers with fluorochrome-conjugated antibodies (Table 5S) in FACS buffer (PBS, 1% FCS, 0.02% Sodium Azide, 2mM EDTA) and 50μl of BD Brilliant stain buffer (BD) for 30 minutes on ice. Cells were washed twice and then fixed and permeabilized with the eBioscience FoxP3/transcription factor staining kit (Thermofisher Scientific). Cells were resuspended in blocking permeabilization buffer (1% mouse serum, 100U/ml heparin, 0.2% BSA) for 20 minutes at room temperature before addition of fluorochrome-conjugated antibodies against intracellular markers (Table S5) on ice for 30 minutes. Cells were washed and resuspended in FACS buffer and stored at 4°C before acquisition on the 5 laser Cytek Aurora instrument (Cytek Biosciences). The data was analyzed on FlowJo LLC software (v10.7.1, BD, Ashland OR)

### Cell sorting for single-cell RNA-sequencing

PBMCs isolated during malaria infection were thawed as described above and stained with Live/dead Fixable Aqua (Thermofisher Scientific) for 30 minutes at room temperature. Cells were washed and incubated with antibodies against CD3, CD4, CD19, IgD, IgM, CD45RO, CD27 and CXCR3 on ice for 30 minutes in MACS buffer (PBS, 2% BSA, 2mM EDTA). Cells were washed, resuspended in a solution of 50% MACS buffer and 50% FBS, and run on the BD Aria III with the 100μm nozzle. Up to 19,000 live naïve CD4^+^ T cells (CD19^-^ CD3^+^ CD4^+^ CD45RO^-^ CD27^+^) or memory CD4^+^ T cells (CD19^-^ CD3^+^ CD4^+^ CD45RO^+^) were sorted into solution of 50% MACS buffer and 50% FBS. Sorted cells were applied to the 10x Genomics single-cell RNA-seq platform for cell barcoding, cDNA synthesis and 5’ gene expression library and V(D)J library preparation following the manufacturer’s protocols. 5’ gene expression and V(D)J libraries were sequenced on the Hi-seq Illumina platform (2x150bp paired-end, Genewiz).

### scRNA-seq analysis and differential expression analysis

Count Matrices for gene expression were generated using the CellRanger software (10x Genomics). After filtering for low quality cells according to the number of RNA, genes detected, and percentage of mitochondrial RNA, dataset were integrated using the standard Seurat v3 integration workflow within each condition, naïve and memory representing a total of 6,664 and 7,934 cells respectively. Integrated data were then used to cluster cells according to their transcriptome similarities. Each cluster was annotated using cell type specific markers.

### Parasite culture and merozoite opsonization assays

*Pf* 3D7 parasites (MR4) were cultured *in vitro* in parasite media (RPMI, 0.6% HEPES, 0.05% gentamicin sulfate, 0.005% hypoxanthine, 0.2% sodium bicarbonate, 0.5% Albumax II, 0.0001% 0.5N NaOH, pH 7.4) with 5% hematocrit in plugged flasks with 1% O2, 5% CO2, 94% N2 gas mixture at 37°C. Cultures were maintained at 2%-4% parasitemia. Merozoite opsonization assay was adapted from previously published protocols ^103, 104^. Briefly, *Pf* 3D7 cultures at ∼2-6% parasitemia were enriched for late stages (trophozoites and schizonts) after diluting the culture by 50% in parasite media and passing it over magnetic LS columns (Miltenyi). Late-stage enriched *Pf* 3D7 cultures were incubated for 12 hours in 10μM E64 (Epoxysuccinyl-L- leucylamido 4-guanidino butane) supplemented parasite media at 37°C with 1% O2, 5% CO2, 94% N2 gas mixture. Late-stage enriched *Pf* 3D7 cultures were then passed through a 1.2μm filter to release merozoites and then passed over a magnetic MS column Fito (Miltenyi) to recover merozoites without hemozoin. Isolated hemozoin-free merozoites were labeled by incubation with 0.5μM YOYO-1 Iodide dye (Thermo Fisher) for 30 minutes at room temperature. Labeled merozoites were washed and centrifuged at 3,700g for 10 minutes before resuspension in RPMI media. Absolute numbers of labeled merozoites were determined by flow cytometry and CountBright™ Absolute Counting Beads (Thermo Fisher). Labeled merozoites were incubated with US control plasma or participant plasma (1:10 dilution or 1:100 dilution) or no plasma for 1 hour at 37°C. Opsonization assay was established with a ratio of 1 THP-1 cell to 10 labeled merozoites pre-incubated with participant plasma (∼2400 THP1 cells: 24,000 labeled merozoites) in FBS blocked 96 well U-bottom plates for 10 minutes at 37°C. Opsonization was terminated by placing cells on ice and washed with cold FACS buffer and centrifuged at 300g for 10 minutes at 4°C. Cells were fixed in 2% PFA for 10 minutes on ice. Cells were washed, resuspended in FACS buffer and stored at 4°C before acquisition on the BD Aria III. The percentage of THP-1 cells positive for YOYO-1 DNA staining was reported.

### Analysis of *Pf* antigen microarray probed with participant plasma

*Plasmodium* arrays were constructed as previously described ^55^. The Pf900/Pf250 array used in the present study, comprising 826 unique features corresponding to 676 unique *Pf* genes, was a down-selected array based on results from previous microarray field studies ^40, 55, 56, 105^. Each microarray chip contained multiple negative *in vitro* transcription/translation control spots that lack plasmid template and serially diluted human immunoglobulin (Ig) G and anti-IgG control spots.

Plasma samples diluted to 1:100 in Protein Array Blocking Buffer (Whatman) were preincubated in 10% of 1mg/ml *Escherichia coli* lysate, and microarrays were probed with plasma samples by incubation overnight at 4°C. The slides were washed 5 times in Tris buffer (pH 7.6) and incubated in Qdot 800-conjugated goat anti–human IgG diluted at 1:100 (Grace Bio-Labs, Inc., Ref # RD480080). The slides were washed 5 times in Tris buffer containing 0.05% Tween 20, followed by a final wash with water. Air-dried slides were scanned at 500ms and quantified using a ArrayCam 400-S Microarray Imaging System (Grace Bio-Labs, Inc.). Quantified signal for each antigen with local background signal correction was considered the raw value. “No- DNA” negative controls consisted *of IVTT* reaction without the addition of plasmid template, representing background *E.coli* antibody levels (no DNA control). The data were normalized by dividing the IVTT protein spot raw intensity by the sample specific mean of the IVTT control spots (no DNA control). This fold-over control (FOC) approach provides a relative measure of the specific antibody binding over the IVTT background. The FOC values were then log2 transformed. A log2 (FOC) value of 0.0 means that the intensity is not different from the background intensity and a value of 1.0 indicates a doubling with respect to the background. A threshold of a log2 (FOC) of 1 was used to define seropositivity. Ags were considered reactive when mean log2 (FOC) among any of the groups is greater than 1.0. Differential analyses of the log2-transfomed data were performed using a Bayes-regularized *t* test for protein arrays. Differences were considered significant with Benjamini Hochberg corrected p-values<0.05. The antibody profile breadth was defined as the number of seropositive Ags (log2 (FOC) > 1) recognized by an individual or for a population using the mean signal intensity for the population. Mann-Whitney p values were calculated by comparing distribution of signals among different groups. Analysis was performed using the R statistical environment (http://www.r-project.org). Graphs were produced in R, Graphpad and Excel software programs.

## Acknowledgments

We thank The Albert Einstein FACS and genomic core facilities (Dave Reynolds), and the Mount Sinai School of Medicine Mass Cytometry core (Adeeb Rahman).

## Author contributions

RF, JPD and GL designed and interpreted most experiments and figures and wrote the paper. KS, AB, AB, ML, TT and JPD contributed to the clinical study design, study management and analysis. PK conducted the sickle cell analysis. FD with RF, JPD and GL analyzed, interpretated all transcriptomic data and contributed to related figures. LL, RF and PF performed protein microarray probing and analysis and contributed to the related figures and tables. JS and SCZ contributed to manuscript preparation. RSK and JS contributed to the biostatistical analysis. RSK was the consulting statistician.

## Data and materials availability

The accession number for the RNA-seq data reported in this paper is GEO:GSE in process. All data is available in the main text or the supplementary materials.

## Supplementary Materials

### Materials and Methods

Figure S1, related to Figure 1.

Figure S2, related to Figure 2.

Figure S3, related to Figure 3 and 4.

Figure S4, related to Figure 5.

Figure S5, related to Figure 6.

Figure S6, related to Figure 7.

Table S1: List of IgG reactive Pf antigens.

Table S2: List of Pf -reactive antigens between patient’s groups.

Table S3: List of IgG-specific Pf antigens with greater reactivity in protected compared to susceptible children.

Table S4: List of antibodies for CyTOF and spectral flow cytometry.

Table S5: List of expressed genes in naive CD4+ T cell clusters defined by single cell transcriptomic.

## Supplementary Figures

**Figure S1.**
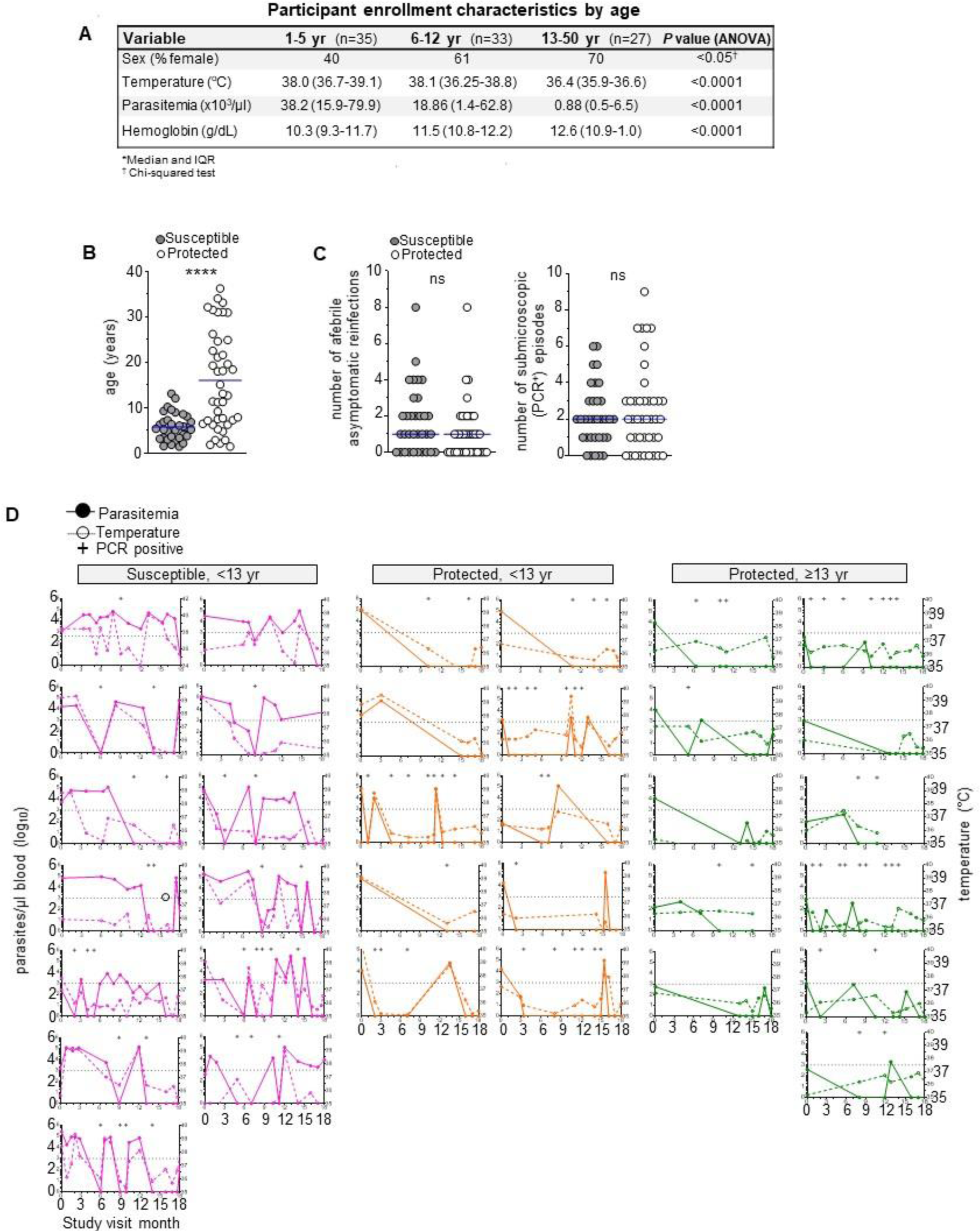
Clinical characteristics of the mild malaria cohort in the 18-month longitudinal study in Malawi. **(A)** Table with participant characteristics and clinical parameters at enrollment in the study, categorized by participants’ age. Median and IQR values, and p-value determined by ANOVA statistical analysis across participants groups is indicated. **(B)** Average number of days between clinical reinfection episodes (>2,500 parasites/µl blood) and age (years) of each participant that completed the study within the protected (≤1 clinical reinfection, n=41) and susceptible groups (≥3 clinical reinfections, n=33). **(C)** Number of afebrile asymptomatic reinfections (<2,500 parasites/µl blood, temperature <37.5°C) and submicroscopic episodes (*Pf* PCR^+^) Student’s t-test was conducted between indicated groups, *p<0.05, **p<0.01, ***p<0.001, ****p<0.0001. **(D)** Parasitemia (left axis), temperature (right axis) and *Pf* PCR status of a subset of susceptible, <13 yr, protected, <13 yr and protected, ≥13 yr at each clinical visit over the 18-month longitudinal study.

**Figure S2.**
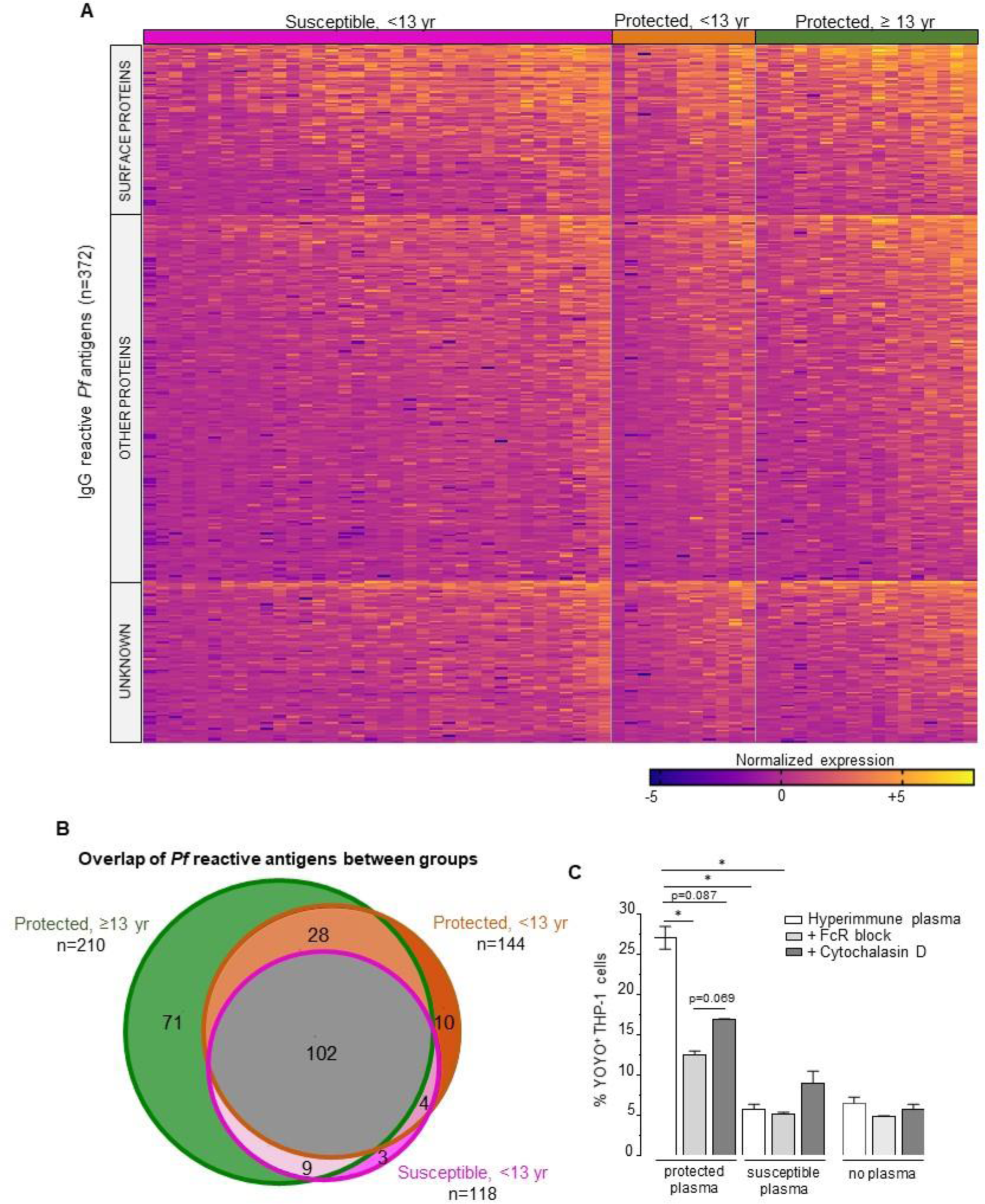
*Pf* specific antibody responses in participants determined by *Pf* antigen array. **(A)** Heatmap of normalized intensity of IgG reactivity to 372 *Pf* Ags (rows) in plasma from individual participants (columns) within susceptible, <13 yr, protected, <13 yr and protected, ≥ 13 yr groups during malaria illness. Surface and non-surface *Pf* antigens are indicated. **(B)** Venn diagram indicating overlapping *Pf* IgG reactivity to 372 *Pf* Ags by susceptible, protected age-matched and protected groups. Total *Pf* Ags recognized by each group is indicated next to the group label. **(C)** *Pf* merozoite opsonization assay with YOYO1^+^ *Pf* merozoites incubated with or without indicated plasma (1:9 dilution, duplicate wells) and co-cultured with THP1 cells in the presence or absence of FcR block or Cytochalasin D. Mean and SEM of the % YOYO1^+^ THP-1 cells are shown. Statistics is Welch’s unpaired t-test between indicated groups *p<0.05.

**Figure S3.**
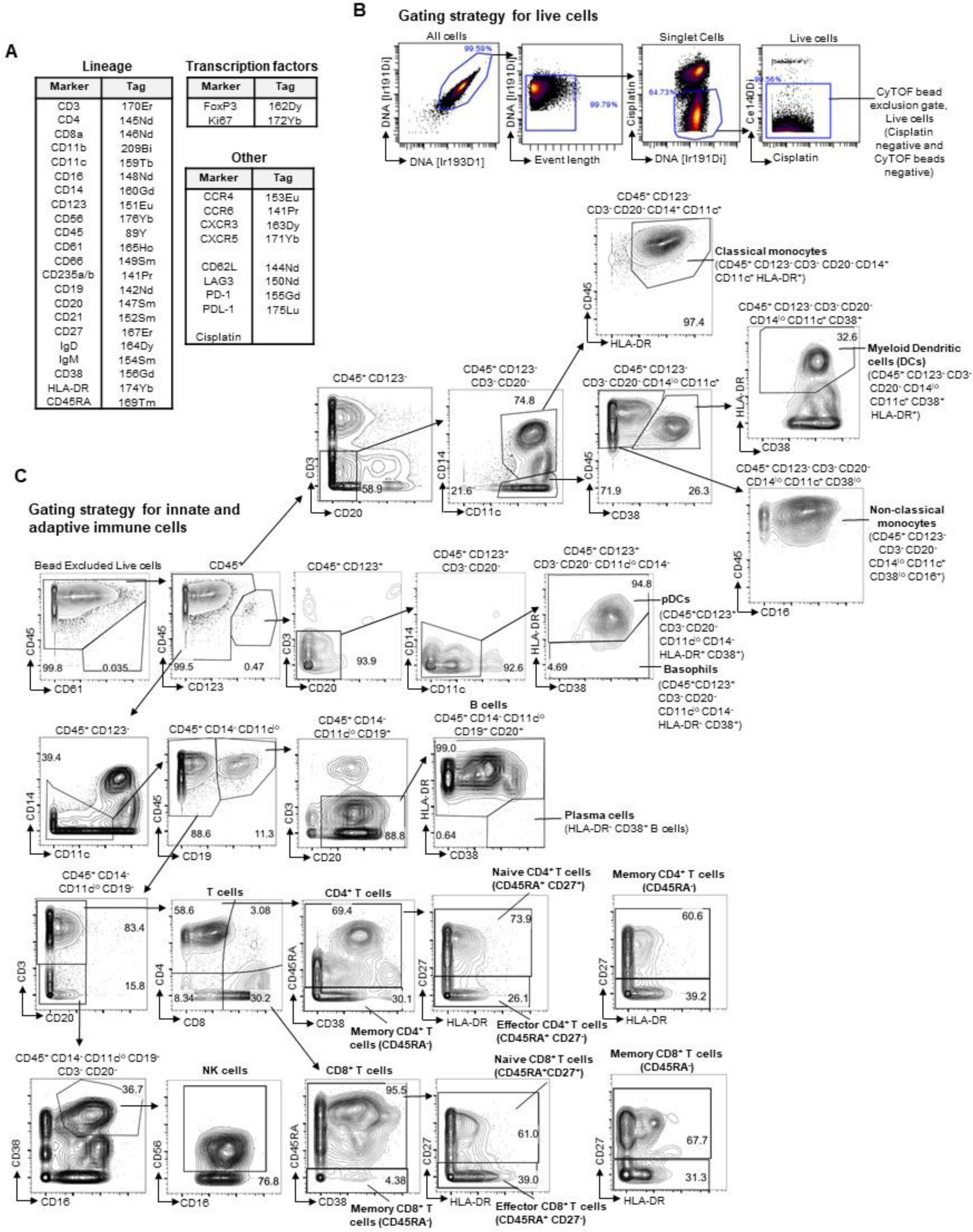
CyTOF immunophenotyping panel and gating strategy of immune cell populations. **(A)** Participant PBMCs were immunophenotyped with a 33-marker mass cytometry antibody panel with indicated heavy-metal tags. **(B)** Gating strategy for live cells among PBMCs immunophenotyped by CyTOF. A density plot that is a representative sample during malaria illness is shown. **(C)** Contour plots with gating strategy for innate and adaptive immune cell populations among PBMCs stained with the antibody panel in (A).

**Figure S4.**
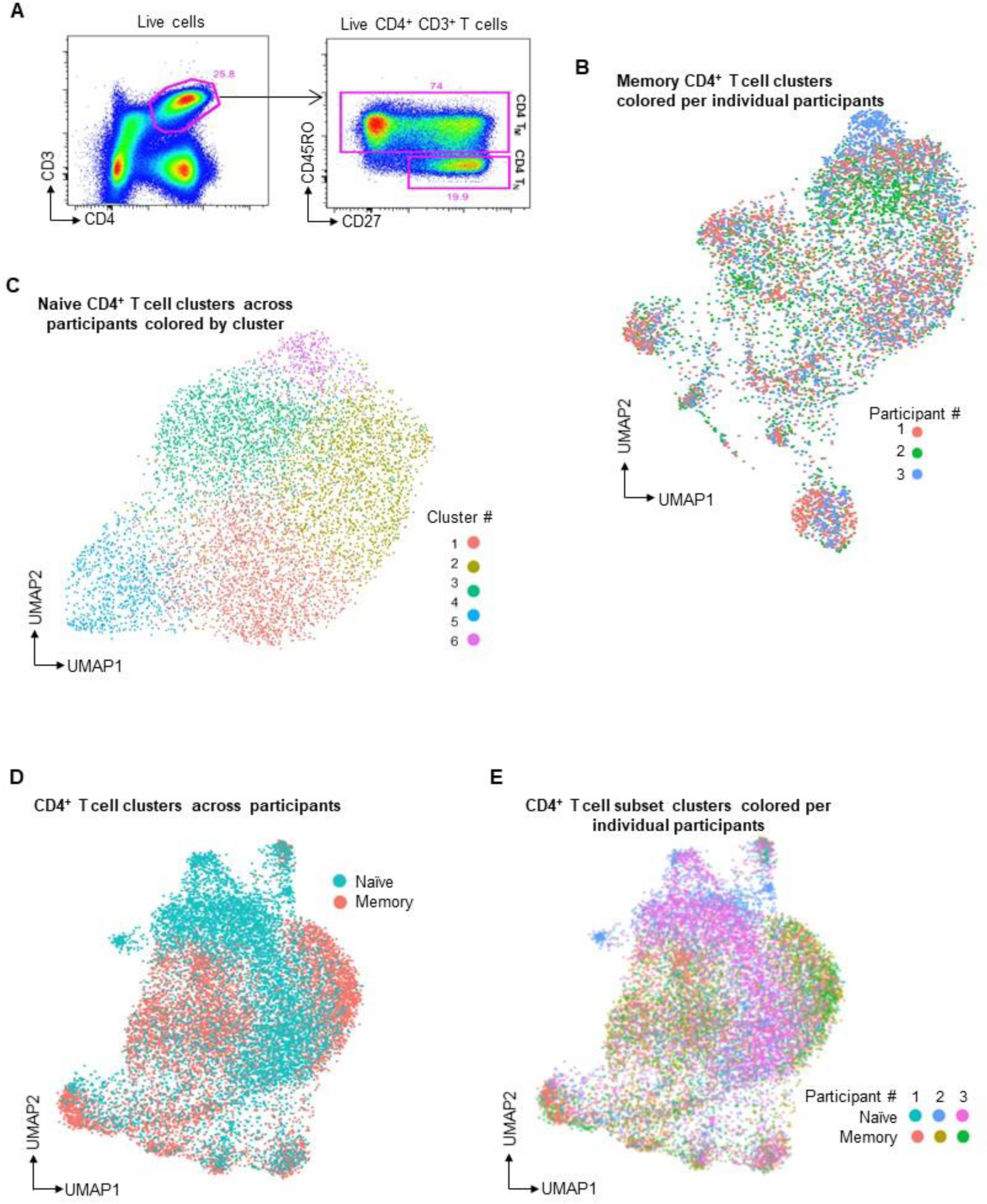
Single-cell transcriptomic analysis of memory and naive CD4^+^ T cells from clinically protected participants during malaria illness. **(A)** Gating strategy for sorting live (L/D aqua negative), memory (CD45RO^+^) and naïve (CD45RO^-^ CD27^+^) CD4^+^ T cells on the BD Aria III for single-cell sequencing analysis. **(B)** UMAP visualization of concatenated single-cell RNA-seq results on isolated memory CD4^+^ T cells from 3 protected participants during malaria infection. **(C)** UMAP visualization of single-cell RNA-seq results on isolated naive CD4^+^ T cells from 3 protected participants during malaria infection. **(D)** UMAP visualization of single-cell RNA-seq results on isolated naive CD4^+^ T cells and memory CD4^+^ T cells from 3 protected participants during malaria infection. **(E)** UMAP visualization of single-cell RNA-seq results on isolated naive CD4^+^ T cells and memory CD4^+^ T cells from 3 protected participants during malaria infection.

**Figure S5.**
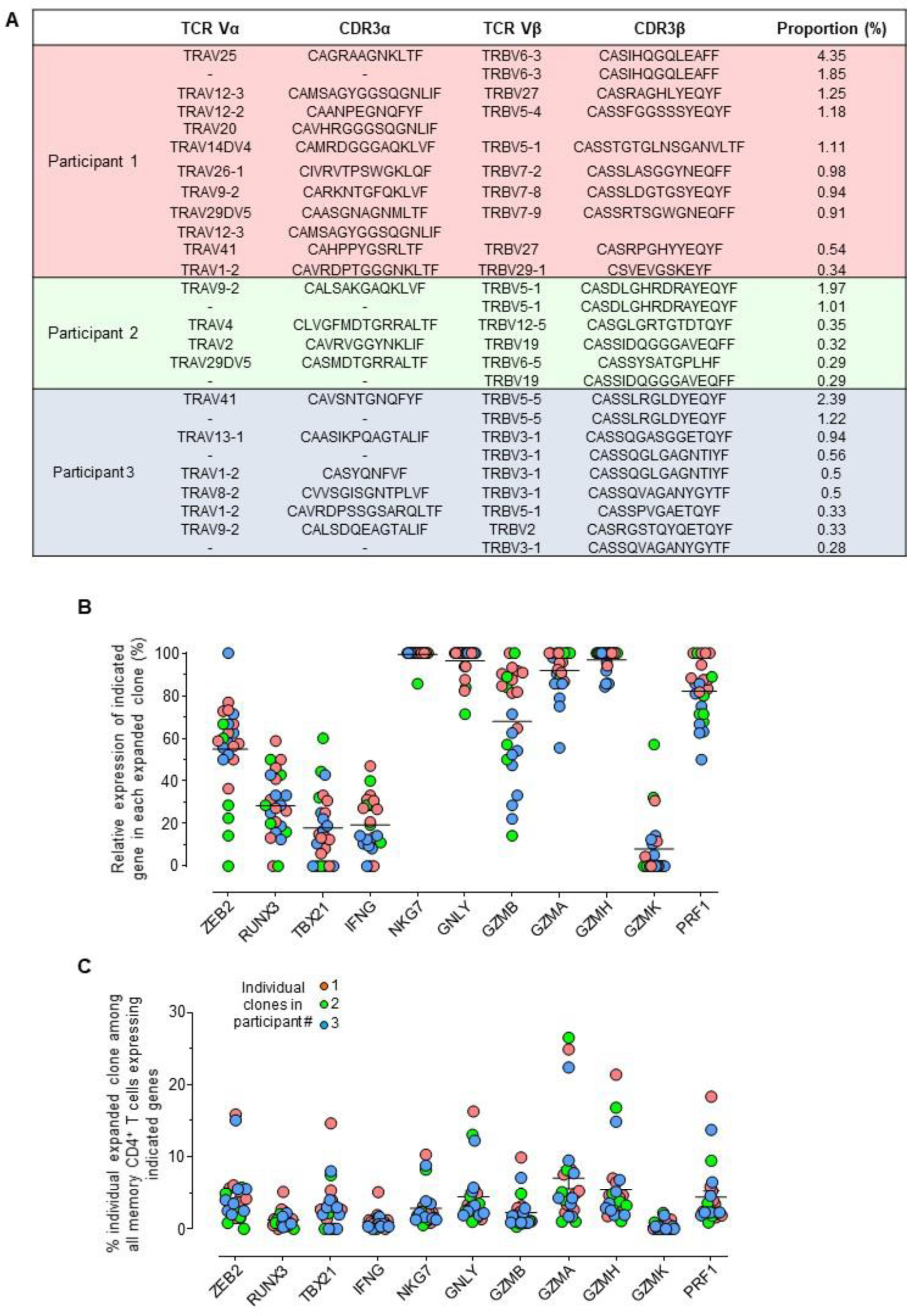
Additional characterization of memory CD4^+^ T cell clonotypes of the 3 clinically protected participants. **(A)** Variable region sequences of the TCRα and TCRβ chains in expanded memory CD4^+^ T cell clones and their frequencies in 3 protected malaria participants, defined by single-cell TCR-seq. **(B)** Relative expression of indicated gene in each individual expanded memory CD4^+^ T cell clone. **(C)** Frequency of expanded memory CD4^+^ T cell clone among all memory CD4^+^ T cells that express the indicated gene in the 3 clinically protected participants.

**Figure S6.**
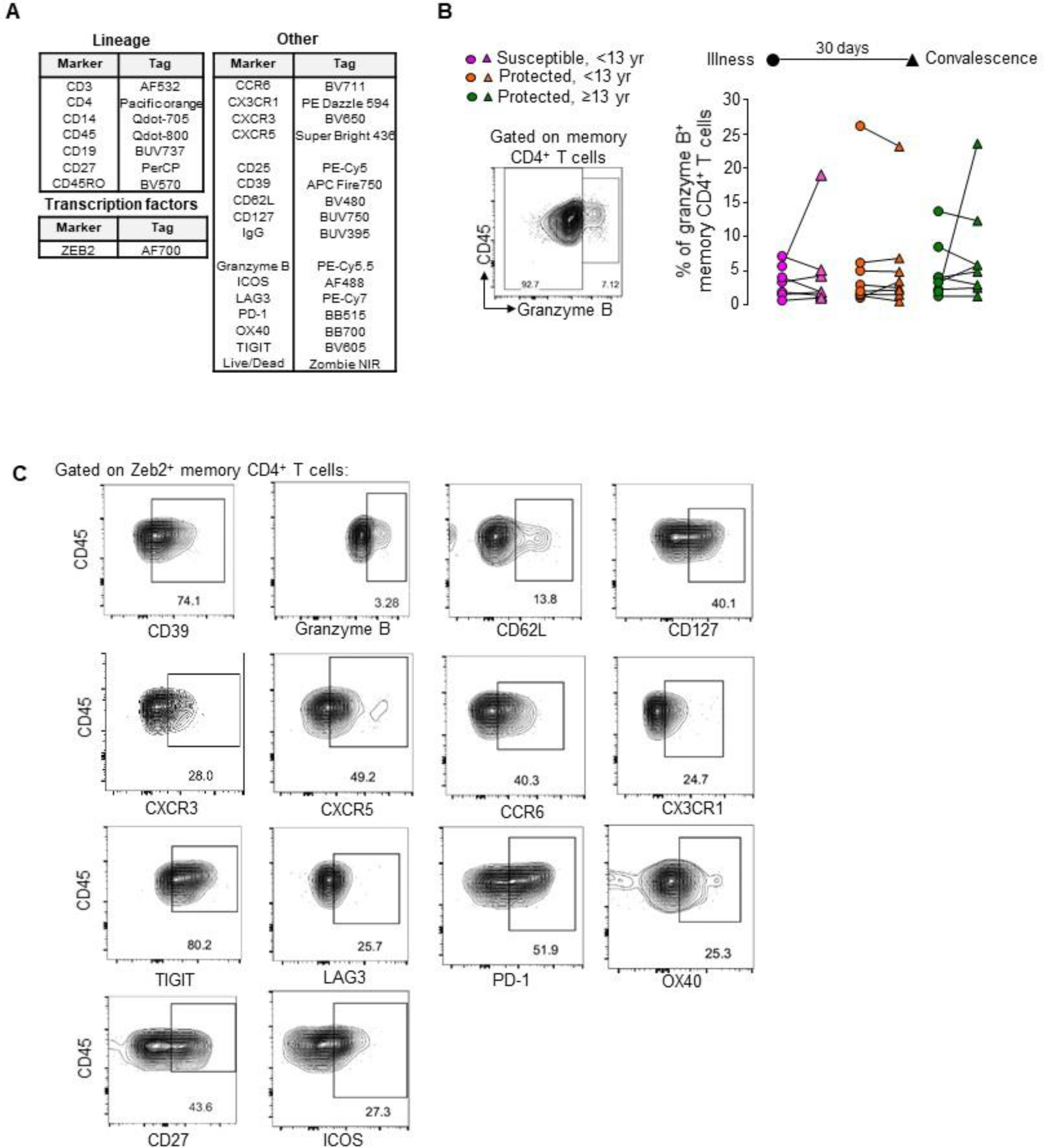
Tracking Zeb2^+^ and Granzyme B^+^ memory CD4^+^ T cells in participants during illness and convalescence. **(A)** Spectral flow cytometry panel (Aurora, Cytek) applied to samples. **(B)** Representative gating of Granzyme B^+^ memory CD4^+^ T cells in a protected patient (left panel). Frequency of Granzyme B^+^ memory CD4^+^ T cells in susceptible, <13 yr (n= 7, pink), protected, < 13 yr (n=8, orange) or protected, ≥13 yr (n=8, green) malaria participants during illness (circle symbol) and at day 30 post illness (convalescence, triangle symbol). Wilcoxon paired t-test for comparisons within each group is shown. **(C)** Representative FACS dot plots of the expression levels of indicated markers on Zeb2^+^ memory CD4^+^ T cells in a representative protected participant. Gates represent the proportion of cell positive for the indicated marker.

